# Diazepam Accelerates GABA_A_R Synaptic Exchange and Alters Intracellular Trafficking

**DOI:** 10.1101/514133

**Authors:** Joshua M. Lorenz-Guertin, Matthew J. Bambino, Sabyasachi Das, Susan T. Weintraub, Tija C. Jacob

## Abstract

Despite 50+ years of clinical use as anxiolytics, anti-convulsants, and sedative/hypnotic agents, the mechanisms underlying benzodiazepine (BZD) tolerance are poorly understood. BZDs potentiate the actions of gamma-aminobutyric acid (GABA), the primary inhibitory neurotransmitter in the adult brain, through positive allosteric modulation of γ2 subunit containing GABA type A receptors (GABA_A_Rs). Here we define key molecular events impacting γ2 GABA_A_R and the inhibitory synapse gephyrin scaffold following initial sustained BZD exposure in vitro and in vivo. Using immunofluorescence and biochemical experiments, we found that cultured cortical neurons treated with the classical BZD, diazepam (DZP), presented no substantial change in surface or synaptic levels of γ2-GABA_A_Rs. In contrast, both γ2 and the postsynaptic scaffolding protein gephyrin showed diminished total protein levels following a single DZP treatment in vitro and in mouse cortical tissue. We further identified DZP treatment enhanced phosphorylation of gephyrin Ser270 and increased generation of gephyrin cleavage products. Selective immunoprecipitation of γ2 from cultured neurons revealed enhanced ubiquitination of this subunit following DZP exposure. To assess novel trafficking responses induced by DZP, we employed a γ2 subunit containing an N terminal fluorogen-activating peptide (FAP) and pH-sensitive green fluorescent protein (γ2^pH^FAP). Live-imaging experiments using γ2^pH^FAP GABA_A_R expressing neurons identified enhanced lysosomal targeting of surface GABA_A_Rs and increased overall accumulation in vesicular compartments in response to DZP. Using fluorescence resonance energy transfer (FRET) measurements between α2 and γ2 subunits within a GABA_A_R in neurons, we identified reductions in synaptic clusters of this subpopulation of surface BZD sensitive receptor. Moreover, we found DZP simultaneously enhanced synaptic exchange of both γ2-GABA_A_Rs and gephyrin using fluorescence recovery after photobleaching (FRAP) techniques. Finally we provide the first proteomic analysis of the BZD sensitive GABA_A_R interactome in DZP vs. vehicle treated mice. Collectively, our results indicate DZP exposure elicits down-regulation of gephyrin scaffolding and BZD sensitive GABA_A_R synaptic availability via multiple dynamic trafficking processes.

## Introduction

GABA_A_Rs are ligand-gated ionotropic chloride (Cl^−^) channels responsible for the majority of fast inhibitory neurotransmission in the adult CNS. The most prevalent synaptic GABA_A_R subtype is composed of two α, two β, and a γ2 subunit forming a heteropentamer (Uusi-Oukari and Korpi, 2010). Benzodiazepines (BZD) are a widely used clinical sedative-hypnotic drug class that selectively binds between the interface of a γ2 subunit and either an α1/2/3/5 subunit (Vinkers and Olivier, 2012). Receptors containing these α subunits are considered to be primarily synaptic, with the exception of α5, which is localized both synaptically and extrasynaptically (Brady and Jacob, 2015). Positive allosteric modulation by BZD enhances GABA_A_R inhibition by increasing the binding affinity of GABA and increasing channel opening frequency (Uusi-Oukari and Korpi, 2010). This potentiating effect of BZD is lost after prolonged or high dose acute exposure (Tietz et al., 1989; Holt et al., 1999), characterized first by a loss of sedative/hypnotic activity followed by the anticonvulsant properties behaviorally (Lister and Nutt, 1986; Wong et al., 1986; File et al., 1988; Bateson, 2002). The induction of BZD tolerance occurs in part due to the uncoupling of allosteric actions between GABA and BZD (Gallager et al., 1984; Marley and Gallager, 1989), a process that appears to rely on GABA_A_R receptor internalization (Ali and Olsen, 2001; Gutierrez et al., 2014). We have previously shown that 24 h BZD treatment leads to decreased surface and total levels of the α2 subunit in cultured hippocampal neurons that was dependent on lysosomal-mediated degradation (Jacob et al., 2012); however, the process by which the α2 subunit is selectively targeted to lysosomes is still unknown. GABA_A_R subunit ubiquitination and subsequent degradation at proteasomes or lysosomes modulates cell surface expression of receptors (Saliba et al., 2007; Arancibia-Carcamo et al., 2009; Crider et al., 2014; Jin et al., 2014; Di et al., 2016). Ubiquitination of the γ2 subunit is the only currently known mechanism identified to target internalized surface GABA_A_Rs to lysosomes (Arancibia-Carcamo et al., 2009).

Another major regulator of GABA_A_R efficacy is postsynaptic scaffolding. Confinement at synaptic sites maintains receptors at GABA axonal release sites for activation. Furthermore, this limits receptor diffusion into the extrasynaptic space where internalization occurs (Bogdanov et al., 2006; Gu et al., 2016). The scaffolding protein gephyrin is the main organizer of GABA_A_R synaptic localization and density, as gephyrin knock-down and knock-out models show dramatic reductions in γ2- and α2-GABA_A_R clustering (Kneussel et al., 1999; Jacob et al., 2005). Evidence suggests gephyrin interacts directly with GABA_A_R α1, α2, α3, α5, β2, and β3 subunits (Kowalczyk et al., 2013; Tyagarajan and Fritschy, 2014; Brady and Jacob, 2015). Gephyrin recruitment is involved in inhibitory long term potentiation (Petrini et al., 2014; Flores et al., 2015), while its dispersal coincides with GABA_A_R diffusion away from synapses (Jacob et al., 2005; Bannai et al., 2009). Extensive post-translational modifications influence gephyrin function (Zacchi et al., 2014; Ghosh et al., 2016). Accordingly, expression of gephyrin phosphorylation site mutants revealed complex effects on GABA_A_R diffusion and gephyrin ultrastructure and scaffolding (Ghosh et al., 2016; Battaglia et al., 2018). Phosphorylation at the gephyrin serine 270 (Ser270) site has been particularly characterized to negatively modulate scaffold clustering and density, in part by enhancing calpain-1 protease mediated degradation of gephyrin (Tyagarajan et al., 2013). Given the well-established interdependent relationship between gephyrin and the γ2 subunit in maintaining receptor synaptic integrity (Essrich et al., 1998; Kneussel et al., 1999; Schweizer et al., 2003; Alldred et al., 2005; Jacob et al., 2005; Li et al., 2005), impaired postsynaptic scaffolding should affect both pre-existing and newly inserted GABA_A_R clustering and ultimately the efficacy of inhibitory neurotransmission. Thus a central unanswered question is if BZD exposure causes changes in gephyrin phosphorylation or protein levels.

Here we demonstrate that 12-24 h treatment with the BZD, diazepam (DZP), leads to a reduction in total γ2 subunit and gephyrin levels in vitro and in vivo. This reduction occurred coincident with enhanced γ2 subunit ubiquitination, but resulted in no significant change in overall γ2 surface levels. Using our recently published dual fluorescent BZD-sensitive GABA_A_R reporter (γ2^pH^FAP), we further show that cell surface γ2-GABA_A_Rs are more frequently targeted to lysosomes after DZP exposure. Forester resonance energy transfer (FRET) experiments further confirmed specific loss of synaptic α2/γ2 GABA_A_R levels following DZP. The scaffolding protein gephyrin also demonstrated augmented phosphorylation at Ser270, increased cleavage and was significantly decreased in membrane and cytosolic compartments. Fluorescence recovery after photobleaching (FRAP) assays identified that DZP treatment increased the simultaneous recovery of γ2-GABA_A_R and gephyrin at synaptic sites, indicating reduced receptor confinement and accelerated exchange between the synaptic and extrasynaptic GABA_A_R pool. This process could be reversed by the BZD site antagonist Ro 15-1788. Lastly, coimmunoprecipitation, quantitative mass spectrometry and bioinformatics analysis revealed shifts in the γ2-GABA_A_R interactome towards trafficking pathways in vivo. Together, these data suggest that DZP exposure causes compensatory decrease in inhibitory neurotransmission by reducing BZD-sensitive GABA_A_R and gephyrin confinement at synapses, and via ubiquitination and lysosomal targeting of γ2.

## Materials and Methods

### Cell Culture, Transfection, Expression Constructs and Mice

Cortical neurons were prepared from embryonic day 18 rats, nucleofected with constructs at plating (Amaxa). The γ2^pH^FAP construct was characterized in (Lorenz-Guertin et al., 2017) and RFP-gephyrin was described in (Brady and Jacob, 2015). The γ2^RFP^ construct was generated by PCR cloning and fully sequenced: the red fluorescent protein mCherry replaced pHluorin in the previously published γ2^pHGFP^ construct (Jacob et al., 2005). GFP-ubiquitin was a gift from Nico Dantuma (Addgene plasmid # 11928) (Dantuma et al., 2006). 8-10 week old male C57BL/6J mice (Jackson Laboratory) were maintained on a reverse 12 h dark/light schedule. Mouse cortical brain tissue was collected and flash frozen 12 h after I.P. injection with either vehicle or diazepam (in 40% PEG, 10% EtOH, 5% Na Benzoate, 1.5 % benzyl alcohol (Hospira)). All procedures were approved by the University of Pittsburgh Institutional Animal Care and Use Committee.

### Reagents, Antibodies, and MG Dye

Diazepam (cell culture, Sigma; injections, Hospira); Ro 15-1788 (Tocris Bioscience); calpain-1 inhibitor MDL-28170 (Santa Cruz); L-glutamic acid (Tocris Bioscience). Primary antibodies: GAPDH (14C10, Cell Signaling); GAD-65 (198104, Synaptic Systems); γ2 GABA_A_R subunit (224003, Synaptic Systems); gephyrin (sc-14003, Santa Cruz); gephyrin (147002, Synaptic Systems); gephyrin mAb7a (147011, Synaptic Systems); GFP (GFP-1020, Aves). MG-BTau dye prepared as in (Lorenz-Guertin et al., 2017).

### Fixed and Live-Imaging

Measurements were made on days in vitro (DIV) 15–19 cortical neurons. Live-imaging performed in Hepes-buffered saline (HBS), containing the following (in mM): 135 NaCl, 4.7 KCl, 10 Hepes, 11 glucose, 1.2 MgCl2, and 2.5 CaCl2 (adjusted to pH 7.4 with NaOH). Images were acquired using a Nikon A1 confocal microscope with a 60× oil objective (N.A., 1.49) at 3× zoom. Data were analyzed in NIS Elements software (Nikon, N.Y.). Measurements were taken from whole cell or averaged from three dendritic 10μm regions of interest (ROI) per cell. For fixed imaging, media was quickly removed and coverslips were washed twice with Dulbecco’s Phosphate Buffered Saline (DPBS) and immediately fixed with 4% paraformaldehyde and then blocked in PBS containing 10% fetal bovine serum and 0.5% bovine serum albumin. Surface antibody staining was performed under non-permeabilized conditions overnight at 4°C. Intracellular staining was performed overnight at 4°C following 0.2% Triton-X permeabilization for 10 min in blocking solution. Synaptic sites were determined during analysis by binary thresholds and colocalization with GAD-65. Extrasynaptic intensity was measured by taking the total dendrite ROI sum intensity minus background and synaptic fluorescence intensity. Dendritic fluorescence was measured using binary thresholds. Experimental conditions were blinded during image acquisition and analysis. The ROUT test (Q=1%) or Grubbs’ Test (alpha=0.05) was used to remove a single outlier from a data set.

### Lysosomal Targeting Assay

Neuron surface and lysosomal-association assays utilized MG-BTau dye for surface receptor pulse-labeling. DIV 15-16 neurons were treated with vehicle or DZP for 8-12 h, then pulse labeled with 100nM MG-BTau for 2 min at room temperature in HBS. Neurons were then washed 5x times with HBS and returned to conditioned media +/− DZP for 1h. To identify lysosomal targeting, 50 nM LysoTracker Blue DND-22 (Life Technologies) and the lysosomal inhibitor, Leupeptin (200μM Amresco), was added 30 min prior to imaging. Following incubation, neurons were washed and imaged in 4°C HBS. Two-three neurons were immediately imaged per culture dish within 10 min of washing. For image analysis, independent ROIs were drawn to capture the soma, three 10 μm sections of dendrite and the whole cell. Binary thresholds and colocalization measurements were performed to identify MG-BTau, pHGFP synaptic GABA_A_R clusters and lysosomes. Total surface pHGFP expression was determined by taking the entire cell surface signal following background subtraction.

### NH_4_Cl Intracellular Imaging

DIV 15-16 neurons were washed and continuously perfused with HBS + treatment at room temperature. Multiposition acquisition was used to image 2-3 neurons per dish. An initial image was taken to identify surface γ2^pH^FAP GABA_A_Rs. Neurons were then perfused with NH_4_Cl solution to collapse the cellular pH gradient and were reimaged. NH_4_Cl solution (in mM): 50 NH_4_Cl, 85 NaCl, 4.7 KCl, 10 Hepes, 11 glucose, 1.2 MgCl_2_, and 2.5 CaCl_2_ (adjusted to pH 7.4 with NaOH). pHGFP intensity was measured following background subtraction and smoothing. Surface/total levels were determined by dividing the first image (surface only) from the second image (total). The spot detection tool in Nikon Elements was used to selectively count larger intracellular vesicles positive for γ2^pH^FAP. A stringent threshold was set to identify brightly fluorescent circular objects with a circumference of approximately 0.75 μm. Values reflect new vesicle objects that were only seen after NH_4_Cl perfusion (second image – first image).

### Intermolecular FRET Imaging, Characterization and Analysis

The α2 pHGFP (α2^pH^) construct was previously published (Tretter et al., 2008) and the γ2^RFP^ construct was generated by PCR cloning and fully sequenced. DIV 15-16 neurons were treated with Veh or DZP for 20-28 h, then washed and continuously perfused with HBS at room temperature. Images were acquired with a 60x objective at 2x zoom. For each cell, an initial image was acquired containing two channels to identify surface α2^pH^ (excited by 488 laser, emission band pass filter 500-550) and γ2^RFP^ participating in FRET (excited 488 FRET, emission band pass filter 575–625 nm, FRET channel). A second, single channel image was taken immediately following with only 561 nm excitation to reveal total γ2^RFP^ levels (excited by 561 laser, emission band pass filter 575–625 nm). For synaptic quantifications, binary thresholding based on intensity was applied with smoothing and size exclusion (0-3 μm) factors. FRET and 561 channel binaries shared identical minimum and maximum binary threshold ranges. Individual synaptic ROIs were created to precisely target and measure synaptic clusters containing both α2^pH^ and γ2^RFP^. Manual trimming and single pixel removal were used to remove signal not meeting the criteria of a receptor cluster. Restriction criteria were applied in the following order: 1) at least 15 synapses measured per cell, 2) FRET γ2^RFP^: raw γ2^RFP^ sum intensity ratio must be less than one, 3) synaptic α2^pH^ mean intensity of at least 500, and 4) α2^pH^ sum intensity limit of 300% of average sum intensity. ROI data was then normalized to vehicle control as percent change. The percentage of RFP participating in FRET was also calculated using FRET RFP:Total RFP ratio.

FRET activity was directly assessed by acceptor (γ2^RFP^) photobleaching. Photobleaching ROIs were implemented on 2 synapses per cell. Pre-bleaching images were acquired every 5 seconds, followed by a γ2^RFP^ photobleaching event using 80% 561 nm laser power. After photobleaching, image capturing resumed without delay using pre-bleach laser power settings for 2 minutes. Image analysis incorporated background subtraction and the measurement of percent change in α2^pH^/FRET γ2^RFP^ ratio over the time course. FRET efficacy measurements compared directly adjacent α2^pH^ and γ2^RFP^ subunits in a GABA_A_R complex. Live-imaging with perfusion of pH 6.0 extracellular imaging saline solution (MES) was used to quench the pH-dependent GFP fluorescence from the α2^pH^ donor fluorophores and show the dependence of FRET on surface α2^pH^ fluorescence. Acidic extracellular saline solution, MES solution pH 6.0 (in mM): 10 MES, 135 NaCl, 4.7 KCl, 11 glucose, 1.2 MgCl_2_, and 2.5 CaCl_2_ (adjusted to pH 7.4 with NaOH). Images were collected under HBS conditions for 1 minute at 20 second intervals, and then followed by a 2 minute MES wash with the same imaging interval to quench donor emissions. FRET RFP mean intensity was measured under both conditions and normalized to HBS. Percent change in FRET RFP emissions were reported.

### Synaptic Exchange Rate FRAP Imaging

Neurons were washed and media was replaced with HBS + treatment. Imaging was performed in an enclosed chamber at 37°C. An initial image was taken for baseline standardization. Photobleaching was performed by creating a stimulation ROI box encompassing two or more dendrites. This stimulation region was photobleached using the 488 and 561 lasers at 25% power for 1 minute. The same stimulation ROI was used for every cell in an experiment. Immediately following photobleaching, 10nM MG-Tau dye was added to the cell culture dish to reidentify surface synaptic GABA_A_R clusters. Time-lapse imaging was then started every 2 min for 60 min. During image analysis, objects were only considered synaptic if they demonstrated colocalization with γ2^pH^FAP pHGFP signal, RFP-gephyrin signal, and had obvious surface MG-BTau fluorescence. ROIs were drawn measuring the rate of fluorescence recovery at 4-8 synaptic sites and one extrasynaptic site (10μm long region; beizer tool) per cell. For data analysis, synapse post-bleach fluorescence intensity time point data was first normalized to pre-bleach fluorescence intensity (post-bleach/pre-bleach). Normalized synapse post-bleach data was then calculated as percent change from t0 ((tx/t0)*100, where × = min). Individual synapses were then averaged to calculate fluorescence recovery and statistically significant changes across time points.

### Western Blot and Immunoprecipitation

Protein concentration was determined by BCA protein assay for all biochemistry. Neurons were lysed in denaturing buffer for immunoprecipitation: 50mM Tris HCl, 1mM EDTA, 1% SDS, 2mM Na_3_VO_4_, 10mM NaF, 50mM N-ethylmaleimide, protease inhibitor cocktail (Sigma). Lysates were sonicated and heated at 50°C for 20 min, then diluted 1:5 in RIPA buffer (50mM Tris HCl pH 7.6, 150mM NaCl, 1% Igepal, 0.5 % Sodium deoxycholate, 1mM EDTA, 2mM Na_3_VO_4_, 10mM NaF, 50mM N-ethylmaleimide, protease inhibitor cocktail). Standard immunoprecipitation procedures were performed using overnight incubation with γ2 subunit antibody or rabbit IgG (sci2027; Sigma), 1 h incubation with protein A sepharose 4B beads (Invitrogen), three RIPA buffer washes, and loading for SDS-PAGE. After electrophoresis and transfer to nitrocellulose membrane, samples were probed with primary antibody overnight followed by the appropriate horseradish peroxide (HRP)-coupled secondary antibody.

### Membrane and Subcellular Fractionation

Cultured neurons were lysed using fractionation buffer: 50mM Tris-HCl, 50mM NaCl, 1mM EDTA, 2mM NaOV, 10mM NaF, 320mM sucrose, 0.25% igepal, and protease inhibitor cocktail. Lysates were spun at 50,000 RPM for 30 min at 4°C to separate pellet (membrane) from supernatant (cytosol). Fraction integrity was tested by localization specific markers in all experiments (Supplementary Figure 1 and data not shown).

### Coimmunoprecipitation

Mice were intraperitoneally (I.P.) injected with vehicle control or 10mg/kg DZP and sacrificed 12 h post-injection (n=4 mice per treatment). Mouse cortical tissue was homogenized in coIP buffer (50mM Tris HCl pH 7.6, 50mM NaCl, 0.25% Igepal, 1mM EDTA, 2mM Na_3_VO_4_, 10mM NaF, 50mM N-ethylmaleimide, and Sigma protease inhibitor cocktail) using a Dounce homogenizer. Tissue was solubilized with end-over-end mixing at 4°C for 15 min, and then spun at 1,000g to remove non-solubilized fractions. Each immunoprecipitation tube contained 375μg of tissue lysate brought up to 1ml volume using coIP buffer. Lysates were precleared using Protein A-Sepharose 4B beads (Invitrogen) for 1 h at 4°C. Lysate was then immunoprecipitated overnight with 2.5μg rabbit γ2 subunit antibody (224003, Synaptic Systems) or 2.5μg rabbit IgG (2027, Santa Cruz). The next day, 40ul Protein A Sepharose slurry was added and mixed for 2 h at 4°C on a nutator. Beads were then washed 3x at 4°C on a nutator in 1ml coIP buffer. Beads were denatured with SDS-PAGE loading buffer [Laemmli Sample buffer (Biorad) + β-mercaptoethanol] with heat at 70°C for 10 min and intermittent vortexing. Two immunoprecipitation reactions were performed per animal and were pooled into a single tube without beads to be used for downstream in-gel digestion.

### Mass Spectrometry and Data Processing

Immunoprecipitated proteins were separated by electrophoresis in Criterion XT MOPS 12% SDS-PAGE reducing gels (Bio-Rad), with subsequent protein visualization by staining with Coomassie blue. Each gel lane was divided into six slices. After de-staining, proteins in the gel slices were reduced with TCEP [tris(2-carboxyethyl)phosphine hydrochloride] and then alkylated with iodoacetamide before digestion with trypsin (Promega). HPLC-electrospray ionization-tandem mass spectrometry (HPLC-ESI-MS/MS) was accomplished by data-dependent acquisiton on Thermo Fisher Orbitrap Fusion Lumos Tribrid mass spectrometer. Mascot (Matrix Science; London, UK) was used to search the MS files against the mouse subset of the UniProt database combined with a database of common contaminants. Subset searching of the Mascot data, determination of probabilities of peptide assignments and protein identifications, were accomplished by Scaffold (v 4.8.4, Proteome Software). MS data files for each entire gel lane were combined via the “MudPIT” option. Identification criteria were: minimum of two peptides; 96% peptide threshold; 1% FDR; 99% protein threshold. One vehicle- and one DZP-treated animal were removed from analysis due to insufficient γ2 subunit pulldown relative to all other groups. Protein clustering was applied in Scaffold and weighted spectrum values and exclusive unique peptides were exported for manual excel analysis. Student’s t-test analysis (P < 0.1) was performed using relative fold change (ratio) of DZP compared to vehicle group. In some cases peptides were only detected in vehicle or DZP treated groups, resulting in DZP/V ratio values of zero or undefined error (cannot divide by zero). These were annotated as NF-DZP (not found in DZP samples) or NF-V (not found in vehicle samples) in the tables.

### Bioinformatics Analysis

Ingenuity Pathways Analysis (IPA) (Ingenuity Systems) was used for cellular pathway analysis. Relative fold levels of DZP proteins compared to vehicle were used for analysis. To be suitable for IPA analysis, proteins NF-DZP were assigned a value of −1E+99, while proteins NF-V were assigned a value of 1E+99. Significant enrichment in protein networks are calculated by right tailed Fisher’s exact test. Z-score analysis is a statistical measure of an expected relationship direction and observed protein/gene expression to predict pathway activation or inhibition. IPA core analysis was searched to determine direct and indirect relationships within 35 molecules per network and 25 networks per analysis. All data repositories available through IPA were used to determine experimentally observed and highly predicted interactions occurring in mammalian tissue and cell lines. Ratio data were converted to fold change values in IPA, where the negative inverse (−1/x) was taken for values between 0 and 1, while ratio values greater than 1 were not affected. Proteins found to be significantly enhanced (P < 0.1) in their association with γ2 were searched in the Mus musculus GO Ontology database (released 2018-10-08) for GO biological process and GO molecular function and analyzed by the PANTHER overrepresentation test; significance was determined using Fisher’s Exact with Bonferroni correction for multiple testing.

## Results

### DZP Exposure Modifies γ2-GABA_A_R and Gephyrin Levels

We first examined if DZP exposure reduced surface levels of γ2-GABA_A_Rs and altered gephyrin S270 phosphorylation in cortical neurons by immunofluorescence (Figure 1A). Cortical neurons were treated for 24 h with vehicle or 1 μM DZP, then immunostained for surface γ2, followed by permeabilization and immunostaining with GAD65 (glutamic acid decarboxylase 65, a marker for presynaptic GABAergic terminals) and the phospho-S270 specific gephyrin mAb7a antibody (Kuhse et al., 2012; Kalbouneh et al., 2014). Image analysis identified no sizable change in surface synaptic (91.6 ± 5.3%) or extrasynaptic (93.3 ± 3.8%) γ2 intensity in DZP treated neurons relative to control, but DZP induced a significant 18.9 ± 7.4% increase in synaptic phospho-gephyrin (Figure 1B). No change in extrasynaptic phosphorylated Ser270 gephyrin was measured. We repeated this DZP treatment and examined total and phospho-gephyrin levels in dendrites (Figure 1C). Again DZP significantly enhanced phospho-S270 gephyrin compared to vehicle (132 ± 12%), while a decrease in overall gephyrin levels was found (69.7 ± 5.4%) (Figure 1D). Accordingly, the mean ratio of phospho/total gephyrin was 78.1± 21% higher following DZP (Figure 1D). Complimentary biochemical studies using membrane fractionation were used to compare cytosolic, membrane, and total protein pools in cortical neurons. In agreement with immunofluorescence data, membrane levels of γ2 (97.1 ± 10.2%) were not reduced after 1 μM DZP, although the total pool of γ2 was diminished (79.3 ± 7.3%) (Figure 2A,B) compared to vehicle. Comparatively, DZP reduced gephyrin in every compartment measured relative to control (cytosol: 87.3 ± 4.6%; membrane: 70.3 ± 10 %; total: 59.1 ± 3.4%). We confirmed the integrity of our fractions using cytosolic and membrane specific markers (Supplemental Figure 1).

**Figure 1.**
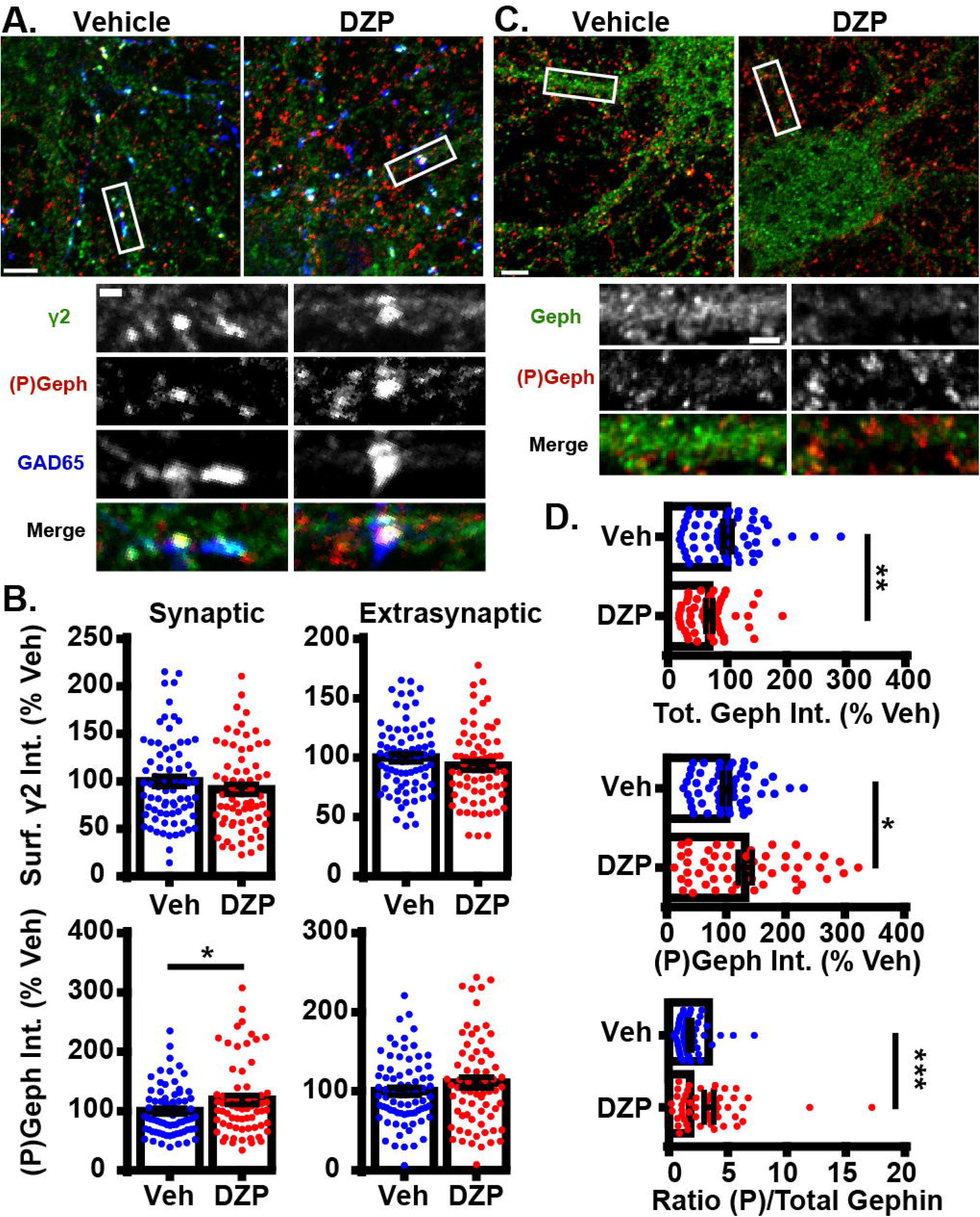
DZP downregulates gephyrin independent of γ2 surface levels. **(A)** Cortical neurons were treated for 24 h with vehicle or 1 μM DZP, then immunostained for surface γ2 GABA_A_R (green), followed by permeabilization and immunostaining for (P)S270 gephyrin (red), and GAD65 (blue). Panels below show enlargements of GABA_A_R synapses. **(B)** Surface synaptic and extrasynaptic γ2 levels are not significantly altered by DZP. Synaptic phospho-gephyrin was enhanced in response to DZP (n=69-74 neurons; 4 independent cultures). **(C)** Neurons were treated as in A followed by antibody staining for total gephyrin (green) and (P)S270 gephyrin (red). Panels below show enlargements of dendrite region. **(D)** The dendritic pool of gephyrin was decreased, while (P)S270 gephyrin levels were augmented, resulting in a dramatic increase in the ratio of phosphorylated gephyrin to total gephyrin (n=52-59 neurons; 3 independent cultures). Image scale bars: main panels = 5 μm, enlargements = 1 μm. *p ≤ 0.05, **p < 0.01, ***p < 0.00, Student’s t-test; error bars ± s.e.m.

**Figure 2.**
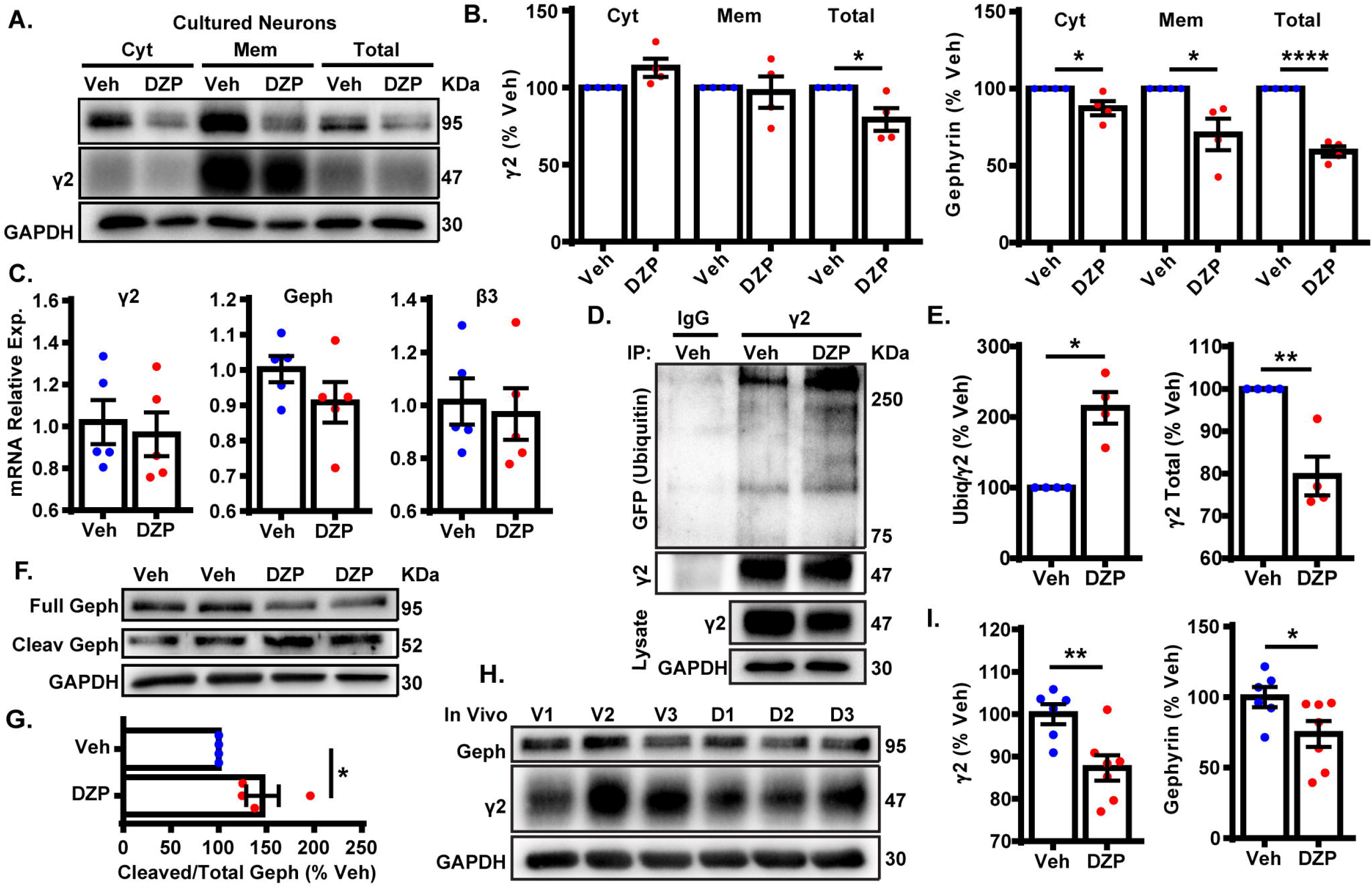
DZP induces degradation of γ2 and gephyrin in vitro and in vivo. **(A)** Cortical neurons exposed to 1 μM DZP or vehicle for 24 h were subjected to membrane fraction and western blot analysis. **(B)** Total γ2 subunit and cytosolic, membrane, and total gephyrin were significantly reduced after DZP (n=4 independent cultures). **(C)** Quantitative RT-PCR revealed no change in γ2 subunit, β3 subunit or gephyrin mRNA expression following 24 h DZP in vitro (n=5 independent cultures). **(D)** GFP-ubiquitin transfected neurons were treated with vehicle or DZP for 12 h. Lysates were immunoprecipitated with control IgG or γ2 antibody, followed by blotting with anti-GFP, γ2 and GAPDH. **(E)** DZP treatment increased the levels of γ2 ubiquitin conjugates and decreased γ2 total levels. **(F,G)** DZP treatment enhanced the ratio of cleaved gephyrin fragments/full length gephyrin (n=4 independent cultures). **(H)** Western blots of cortical tissue collected from mice 12 h after a single IP injection of 10 mg/kg DZP or vehicle. **(I)** γ2 subunit and gephyrin totals are significantly reduced in DZP-treated animals (6-7 mice per condition). (*p ≤ 0.05, **p < 0.01, ****p < 0.0001, Student’s t-test; error bars ± s.e. m)

Next we assessed if the decrease in gephyrin and γ2 total levels at 24 hours was a result of altered gene expression. qRT-PCR experiments revealed no difference in gephyrin, γ2, or control GABA_A_R β3 subunit mRNA levels between vehicle and DZP treated neurons (Figure 2C). To determine if post-translational modification of γ2 also occurs coincident with decreased γ2 protein levels, we examined ubiquitination of γ2 in response to DZP exposure. We reasoned that changes in ubiquitination of γ2 would likely precede the loss of total γ2 seen at 24 h (Figure 2A,B). GFP-ubiquitin transfected cortical neurons were treated with vehicle or 1 μM DZP for 12 h. Neurons were lysed under denaturing conditions to isolate the γ2 subunit from the receptor complex (Supplemental Fig. 2). Immunoprecipitation of the γ2 subunit revealed a 2.13 fold increase in ubiquitination in DZP treated neurons relative to vehicle (Figure 2D,E). Furthermore, just as observed with 24 h DZP treatment, a reduced total pool of γ2 was also found at 12 h (Figure 2D,E). Notably, this is the first demonstration of endogenous γ2 ubiquitination occurring in neurons (previous findings were of recombinant receptors in HEK cells) (Arancibia-Carcamo et al., 2009; Jin et al., 2014). To investigate mechanisms underlying reduced gephyrin levels, we examined gephyrin cleavage. Gephyrin is degraded post-translationally by the protease calpain-1 (Tyagarajan et al., 2013; Costa et al., 2015; Kumar et al., 2017), and gephyrin Ser270 phosphorylation promotes cleavage by calpain-1 (Tyagarajan et al., 2013). Consistent with the enhanced gephyrin S270 phosphorylation (Figure 1) and reduced total levels (Fig. 1,2) we found a significant increase in the ratio of cleaved/full length gephyrin after 24 h DZP in vitro (Fig. 2F,G). We confirmed the identity of the gephyrin cleavage product using a well-characterized glutamate stimulation protocol that induces gephyrin cleavage in cultured neurons (Costa et al., 2015; Kumar et al., 2017), a process blocked by calpain-1 inhibition (Supplemental Figure 3).

Finally, we wanted to determine if similar mechanisms occur in vivo following DZP treatment. Prior publications show that BZDs and metabolites are not present 24 h post-injection due to rapid drug metabolism in rodents (Yoong et al., 1986; Xie and Tietz, 1992; Van Sickle et al., 2004; Markowitz et al., 2010). Furthermore, BZD uncoupling does not persist 24 h after a single dose (15 mg/kg) or 2 week daily DZP treatment, whereas uncoupling can be seen 12 h after a single injection, indicating this is the appropriate timepoint for measuring in vivo loss of γ2-GABA_A_R function (Holt et al., 1999). Accordingly, mice were given a single intraperitoneal (IP) injection of 10 mg/kg DZP or vehicle control, and cortex tissues were harvested 12 h later. We found DZP significantly reduced the total pool of γ2 (87.3 ± 3.0%) and gephyrin (73.9 ± 9.1%) relative to vehicle treated mice at 12 h post injection (Figure 2H,I). These findings indicate both BZD-sensitive GABA_A_Rs and gephyrin are downregulated by post-translational mechanisms after initial DZP treatment in vitro and in vivo to temper potentiation of GABA_A_R function.

### DZP Enhances Intracellular Accumulation and Lysosomal Targeting of γ2-GABA_A_Rs

We then investigated if surface DZP-sensitive GABA_A_Rs are more frequently targeted to lysosomes after DZP exposure by live-imaging. For these experiments we used our recently characterized optical sensor for synaptic GABA_A_R (γ2^pH^FAP). This dual reporter is composed of a γ2 subunit tagged with an N terminal pH-sensitive GFP, myc, and the fluorogen-activating peptide DL5 (Lorenz-Guertin et al., 2017). The pH-sensitive GFP tag selectively identifies cell surface GABA_A_Rs and the DL5 FAP binds malachite green (MG) dye derivatives including MG-BTau (Szent-Gyorgyi et al., 2008; Szent-Gyorgyi et al., 2013; Pratt et al., 2017). MG-BTau is cell impermeable and non-fluorescent until bound by DL5. Upon binding, MG-BTau fluoresces in the far red spectral region (~670 nM). This FAP-dye system allows for selective labeling of surface γ2-containing GABA_A_Rs which can then be tracked through various phases of trafficking (Lorenz-Guertin et al., 2017). As previously shown, γ2^pH^FAP GABA_A_Rs are expressed on the neuronal surface, form synaptic clusters, do not perturb neuronal development and show equivalent functional responsiveness to GABA and DZP both in the absence and presence of MG dyes (Lorenz-Guertin et al., 2017). We transfected neurons with γ2^pH^FAP and treated them with DZP for 8-16 h. Neurons were then pulse-labeled with 100 nM MG-BTau dye and returned to conditioned media at 37°C +/− DZP for 1 h. The lysosomal inhibitor leupeptin (200 μM) and the lysosomal specific dye, Lysotracker (50 nM), were added after 30 min. At the end of the incubation, neurons were washed in 4°C saline to inhibit trafficking and immediately used for live-imaging experiments. Representative images demonstrate MG-BTau labeled γ2^pH^FAP-GABA_A_Rs localized on the cell surface (Figure 3A) and at synaptic clusters on dendrites (Figure 3B) based on colocalization with surface specific pHGFP signal. MG-BTau further reveals internalized receptors at lysosomes (Figure 3C). Image quantification showed synaptic γ2-GABA_A_R intensity remained largely unchanged (Figure 3D). Importantly, we found a significant 8.0 ± 2.5% enhancement in the mean intensity of GABA_A_Rs labeled with MG-BTau at lysosomes following DZP (Figure 3E).

**Figure 3.**
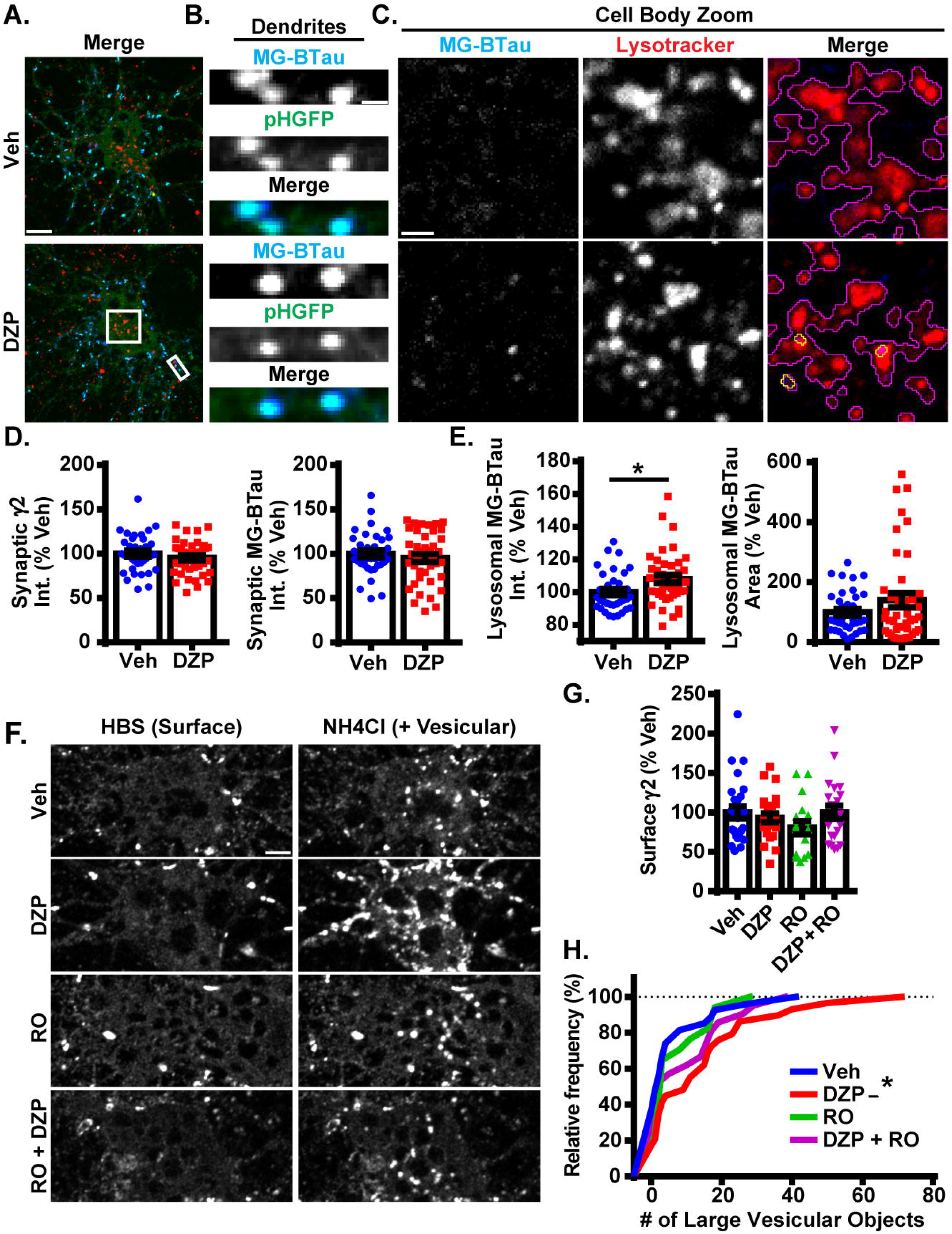
Lysosomal targeting and vesicular accumulation of γ2-GABA_A_Rs in response to DZP. **(A)** γ2^pH^FAP neurons were pretreated for 12-18 h with 1 μM DZP, then pulse-labeled with 100 nM MG-BTau dye for 2 min, and returned to conditioned media at 37°C +/− DZP for 1 h. 50 nM Lysotracker dye was added at the 30 min mark to identify lysosomes. MG-BTau = blue; pHGFP = green; Lysotracker = red. (n=37-42 neurons; 5 independent cultures). **(B)** Dendrite zoom images show MG-BTau labeling at γ2^pH^FAP synapses. **(C)** Cell body zoom images highlighting colocalization of MG-BTau labeled GABA_A_Rs (yellow trace) at lysosomes (purple trace). **(D)** pHGFP and MG-BTau measurements reveal surface synaptic γ2-GABA_A_R levels are not altered by DZP. **(E)** The pool of internalized MG-BTau GABA_A_Rs colocalized at lysosomes was enhanced in DZP treated neurons as measured by intensity (*p ≥ 0.05, Student’s t-test; error bars ± s.e.m). **(F)** Neurons treated 20-28 h with vehicle or DZP. The DZP site antagonist Ro 15-1788 (5 μM) was added 1-2 h prior to imaging to inhibit DZP binding at GABA_A_Rs. Neurons were first imaged in HBS, and then perfused with NH_4_Cl (pH 7.4) to reveal intracellular γ2^pH^FAP receptors. DZP treated neurons accumulated more γ2-GABA_A_Rs in large vesicular structures compared to vehicle. (n=20-27 neurons; 3-4 independent cultures). **(G)** Surface intensity of γ2^pH^FAP was not different between treatments (one-way ANOVA; error bars ± s.e.m). **(H)** DZP-treated neurons more frequently demonstrated accumulation of γ2^pH^FAP in large vesicles (*p ≤ 0.05 Kolmogorov-Smirnov statistical test). Scale bars in μm: A=10; B=1;C=2, F=5.

We complemented these lysosomal targeting studies with an NH_4_Cl live-imaging approach that allows us to compare the ratio of cell surface vs. intracellular GABA_A_Rs in living neurons. γ2^pH^FAP expressing neurons were treated with vehicle or DZP for 24 h. Additional control groups included the BZD antagonist Ro 15-1788 (1-2 h) to reverse the effects of DZP. Neurons were actively perfused with HEPES buffered saline (HBS) treatment and an initial image was taken of surface pHGFP receptor signal (Figure 3F). Neurons were then exposed to pH 7.4 NH4Cl solution to neutralize the pH gradient of all intracellular membrane compartments, revealing internal pools of γ2 containing GABA_A_Rs. Analysis revealed no change in surface γ2 levels between treatments (Fig. 3G) consistent with (Figures 1, 2). However, the number of large intracellular vesicles (circular area ~0.75 μm) containing receptors was significantly enhanced (Figure 3H), consistent with increased localization in intracellular vesicles. Ro 15-1788 and DZP + Ro 15-1788 treated neurons were not significantly different from vehicle. Overall, these findings suggest γ2-GABA_A_R ubiquitination, intracellular accumulation, lysosomal targeting and degradation are part of the adaptive response to DZP.

### Surface Levels of Synaptic α2/γ2 GABA_A_R are Decreased Following DZP

Despite the increase in ubiquitination and lysosomal targeting of γ2-GABA_A_Rs after DZP, we did not detect decreased overall surface or synaptically localized surface γ2 levels. This suggested two possibilities, one being that DZP treatment only reduced total γ2 levels to 80% of control in cultured cortical neurons and 85% in vivo, making a slight decrease in surface γ2-GABA_A_Rs challenging to detect with current methods. Alternatively, there could be an increase in γ2 subunit assembly with BZD-insensitive α subunits (γ2α4β) (Wafford et al., 1996) with a concomitant reduction in surface levels of BZD-sensitive receptors (γ2α1/2/3/5β). Our previous work showed 24 h BZD exposure in hippocampal neurons causes decreased total and surface levels of the α2 GABA_A_R subunit via lysosomal mediated degradation, without any changes in receptor insertion or removal rate (Jacob et al., 2012). To determine if α2/γ2 GABA_A_Rs are specifically decreased by DZP treatment, we developed and employed an intermolecular FRET assay, using pH-sensitive GFP tagged α2^pH^ (Tretter et al., 2008) as a donor fluorophore and a red fluorescent protein (RFP) tagged γ2 subunit (γ2^RFP^) as an acceptor. FRET is an accurate measurement of molecular proximity at distances of 10-100 Å and is highly efficient if donor and acceptor are within the Förster radius, typically 30-60 Å (3-6 nM), with the efficiency of FRET being dependent on the inverse sixth power of intermolecular separation (Förster, 1965). Synaptic GABA_A_Rs exist as five subunits assembled in γ2-α-β-α-β order forming a heteropentamerc ion channel (Figure 4A). We first expressed α2^pH^ and γ2^RFP^ in neurons and examined their ability to participate in intermolecular FRET. Photobleaching of the acceptor γ2^RFP^ channel enhanced donor α2^pH^ signal (Supplementary Figure 4), confirming energy transfer from α2^pH^ to γ2^RFP^. Next, we confirmed measurable FRET only occurs between α2^pH^/γ2^RFP^ in surface GABA_A_R at synaptic sites; FRET was blocked with quenching of donor α2^pH^ when the extracellular pH was reduced from 7.4 to 6.0 (Figure 4A,B). Following FRET assay validation, α2^pH^/γ2^RFP^ GABA_A_R expressing neurons were treated for 24 h with vehicle or DZP and examined for total synaptic α2^pH^ and γ2^RFP^ fluorescence as well as the γ2 FRET signal (Fig. 4A). These studies identified a DZP-induced reduction in synaptic α2 (−12.6%), synaptic γ2 (−14.3%) and diminished association of α2 with γ2 in synaptic GABA_A_Rs as measured by decreased FRET γ2 signal (−10.6%) (Figure 4B). In summary, this sensitive FRET method indicates that cortical neurons show a similar susceptibility for α2 subunit downregulation by BZD treatment as seen in hippocampal neurons (Jacob et al., 2012). Furthermore it identifies a DZP-induced decrease in a specific pool of surface synaptic BZD-sensitive γ2-GABA_A_R.

**Figure 4.**
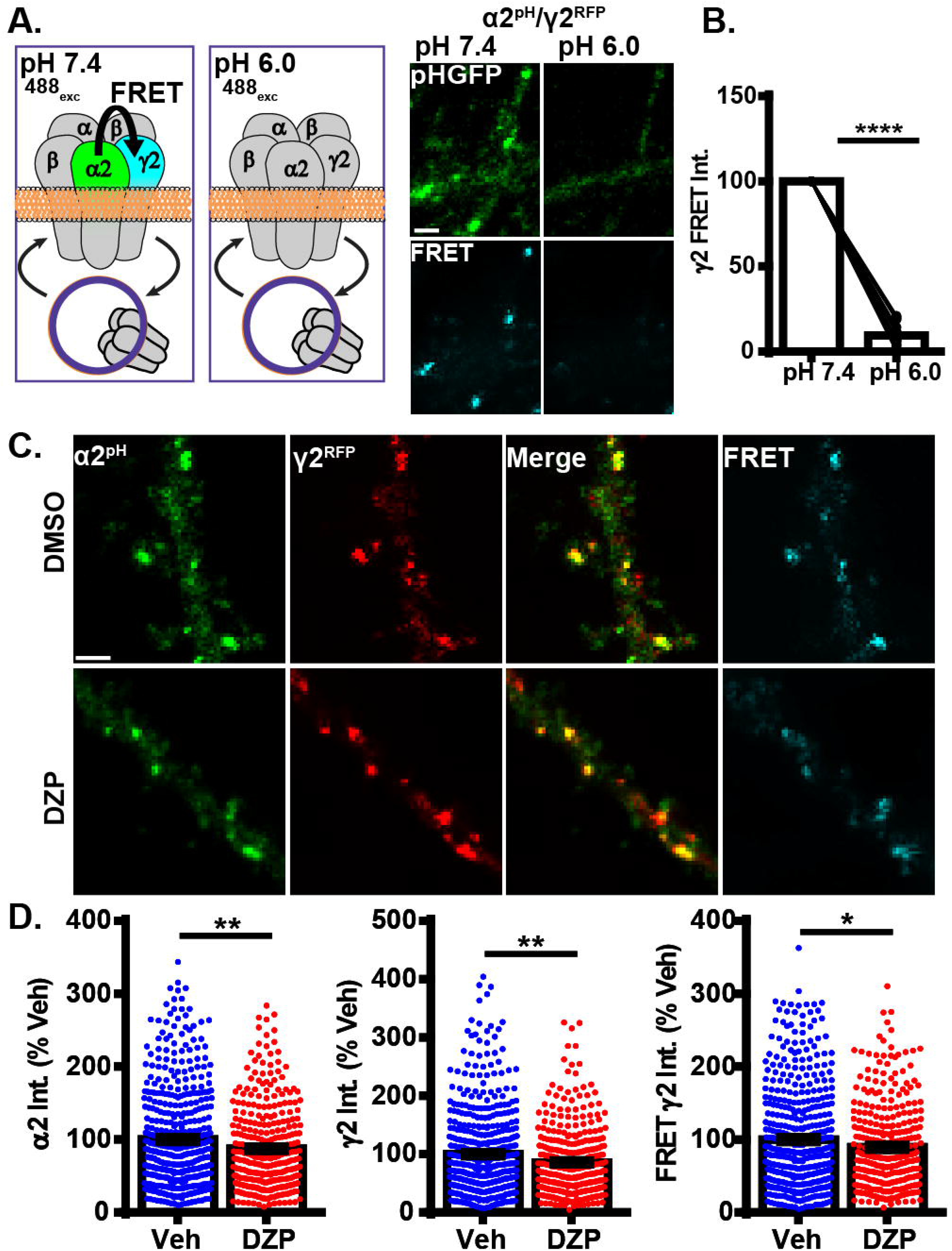
Intermolecular FRET reveals decreased synaptic α2/γ2 surface GABA_A_Rs after DZP. **(A)** Diagram and time-series images of cortical neurons expressing donor α2^pH^ (green) and acceptor γ2^RFP^ during imaging and with imaging saline at pH 7.4 and pH 6.0. Surface α2^pH^ (green) signal and intermolecular FRET (teal) between α2/γ2 subunits occurs at pH 7.4, but is eliminated by brief wash with pH 6.0 extracellular saline and quenching of the α2^pH^ donor pHGFP fluorescence. **(B)** Quantification of relative FRET at pH 7.4 and pH 6.0 (n = 20 synapses). **(C)** Neurons α2^pH^ (green) and γ2^RFP^ (red) were treated with vehicle or DZP for 20-28 h +/− and then subjected to live-imaging. For each cell, an initial image used 488 nm laser excitation to identify surface α2^pH^ and FRET γ2^RFP^. A second image was taken immediately afterwards to acquire γ2^RFP^ total levels (561 nm laser excitation). Dendritic lengths show multiple synaptic clusters with α2/γ2 surface GABA_A_Rs. **(D)** Synaptic cluster intensity quantification of α2^pH^, γ2^RFP^ and FRET γ2^RFP^ (at least 15 synapses per cell; n = 335-483 synapses; 6 independent cultures). Image scale bars = 2 μm (*p ≤ 0.05, **p < 0.01, ****p < 0.0001, paired t-test (B), Student’s t-test (D); error bars ± s.e.m).

### Synaptic Exchange of γ2-GABA_A_Rs and Gephyrin are Accelerated after Prolonged DZP Treatment

We previously found 24 h BZD exposure reduces the amplitude of miniature inhibitory postsynaptic currents (mIPSC) (Jacob et al., 2012), suggesting changes in synaptic GABA_A_R function. Having identified both reductions in gephyrin (Figures 1,2) and BZD sensitive GABA_A_Rs (Figures 2, 4), we next tested if DZP treatment altered the synaptic retention properties of gephyrin and/or GABA_A_Rs. Neurons expressing γ2^pH^FAP and RFP-gephyrin were used for live-imaging fluorescence recovery after photobleaching experiments (FRAP) to measure synaptic and extrasynaptic exchange following exposure to vehicle, 1 μm DZP, 5 μm Ro 15-1788, or DZP + Ro 15-1788. After an initial image was taken, dendrites were photobleached, and signal recovery was measured every 2 min over 30 min at synaptic sites and extrasynaptic regions (Figure 5A synapses panel; Figure 5B larger dendritic region with asterisk denoting extrasynaptic region). MG-BTau dye was added directly after the photobleaching step to immediately re-identify the photobleached surface synaptic GABA_A_Rs, and improve spatial measurements (Figure 5B). These experiments revealed synaptic γ2 turnover rates were nearly doubled in DZP treated neurons, a process reversed by Ro 15-1788 co-treatment (Figure 5C). DZP also accelerated gephyrin synaptic exchange rates compared to vehicle, with Ro 15-1788 co-treatment restoring exchange to control levels. No significant correlation was found between cluster area measured and fluorescence recovery rates of γ2 and gephyrin across all conditions, suggesting synaptic exchange rate is independent of cluster size (Supplementary Figure 5). Moreover, no statistical difference was found in γ2 or gephyrin extrasynaptic exchange rates (Figure 5D). These findings suggest concurrent reduction of gephyrin and GABA_A_R synaptic confinement is a compensatory response to mitigate prolonged DZP potentiation of GABA_A_RS.

**Figure 5.**
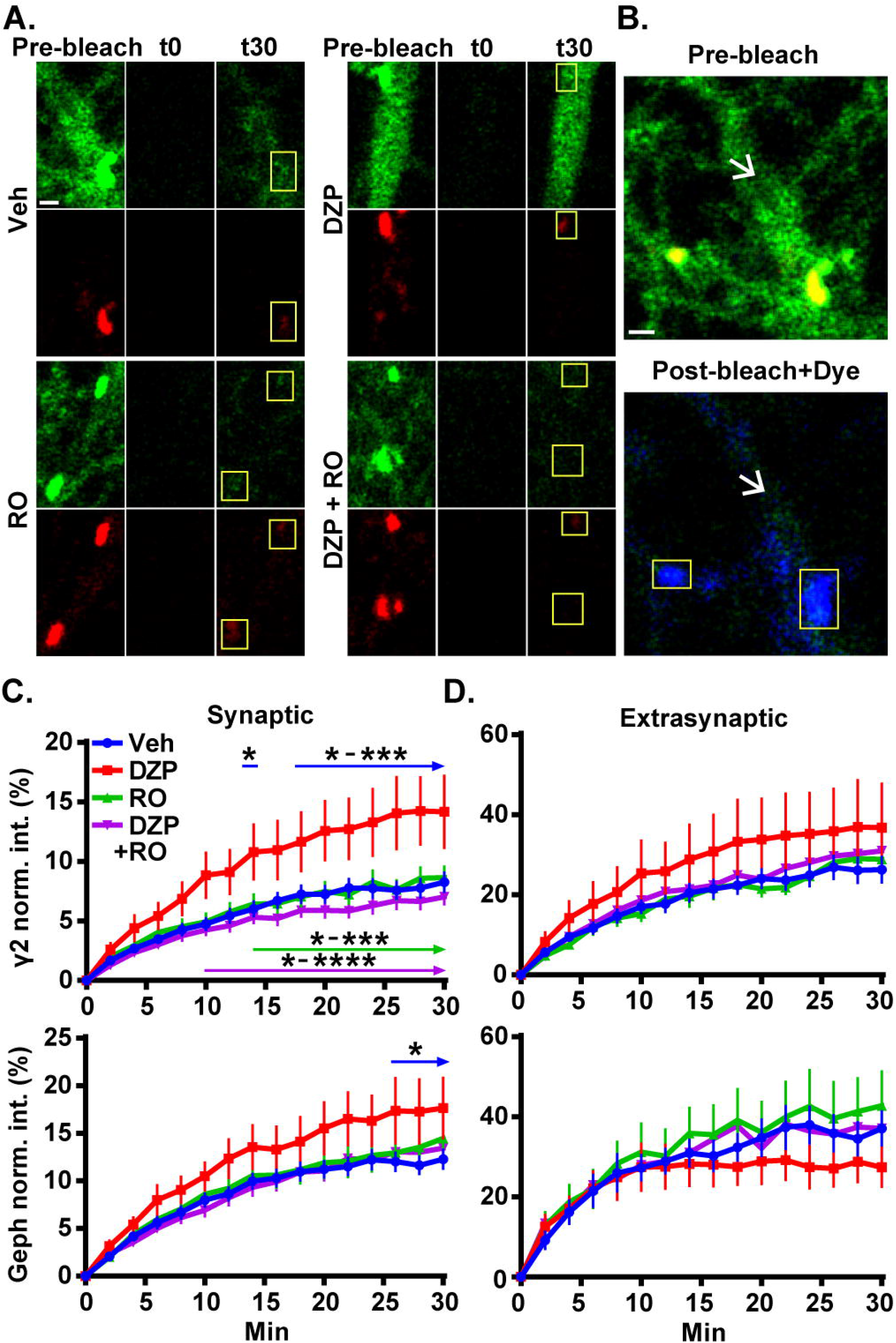
Prolonged DZP exposure accelerates γ2 GABA_A_R and gephyrin synaptic exchange. **(A)** Neurons expressing γ2^pH^FAP GABA_A_R (green) and RFP-gephyrin (red) were treated with vehicle or DZP for 20-28 h +/− Ro 15-1788 for the last 1-2 h. Neurons were imaged at 37°C in constant presence of treatment. Initial image taken prior to photobleaching (Pre-bleach), then imaged postbleach (t0) every 2 min for 30 min. Images of dendritic regions show synaptic cluster sites (yellow boxes) and extrasynaptic regions. **(B)** 10 nM MG-BTau dye (blue) was added immediately after bleaching events in A. to resolve bleached γ2^pH^FAP GABA_A_Rs and provide spatial accuracy for time series measurements. Yellow box indicates synaptic cluster site, and arrows indicates extrasynaptic region. **(C, D)** Fluorescence recovery of γ2 GABA_A_R and gephyrin measured at synaptic sites and extrasynaptic sites from A. Synapse = γ2^pH^FAP cluster colocalized with gephyrin cluster. Image scale bars = 1 μm (*p ≤ 0.05, **p < 0.01, ***p < 0.001, ****p < 0.0001, two-way ANOVA; Tukey’s multiple comparisons test; 4-8 synapses and one 10 μm extrasynaptic region per cell; 51-56 synapses from 16 neurons per treatment; 4 independent cultures; error bars ± s.e. m).

### Coimmunoprecipitation and Quantitative Proteomics of γ2 GABA_A_R Following DZP Injection

We sought to observe DZP-induced changes in receptor trafficking in vivo. As an orthogonal approach, we utilized label-free quantitative proteomics to measure changes in the quantities of proteins associated with γ2-GABA_A_Rs in the cortex of mice after DZP. Cortical tissue was collected from DZP-or vehicle-treated mice 12 h post injection, lysed, and immunoprecipitated with anti-γ2 subunit antibody or IgG control. Following label-free mass spectrometry analysis, spectrum counts were used to assess relative abundance of γ2-associated proteins. A total of 395 proteins was identified using our inclusion criteria: minimum of two peptides; identified in at least three samples overall or in two of three samples in a specific treatment group; demonstrated at least 3:1 enrichment over IgG control across at least three samples overall (Dataset 1). The relative abundance of γ2-GABA_A_R associated proteins in the DZP group compared to vehicle was used to determine which proteins were significantly (P < 0.1) increased (Table 1) or decreased (Table 2). As a result we identified 46 proteins with elevated levels of interaction with γ2-GABA_A_Rs, including 10 proteins that were only found in the DZP treated group (Table 1, not found in vehicle samples, NF-V). Notably, we found a significant increase in γ2 association with 14-3-3 protein family members, the heat shock protein family A (Hsp70) member 8 (also known as Hsc70) and the GABA_A_R α5 subunit, suggesting DZP induced changes in GABA_A_R surface trafficking (Qian et al., 2012; Nakamura et al., 2016a), gephyrin clustering (Machado et al., 2011) and receptor composition (van Rijnsoever et al., 2004). In contrast, 23 proteins were found to coimmunoprecipitate with γ2 less in DZP animals relative to control, seven of which were only present in the vehicle treatment group (Table 2, not found in DZP, NF-DZP). Interestingly, the calcium-sensitive kinase CaMKIIα, which can regulate GABA_A_R membrane insertion, synaptic retention and drug binding properties (26, 70-72), was found to be significantly decreased in interaction with γ2-GABA_A_R following DZP injection in vivo.

**Table 1.**
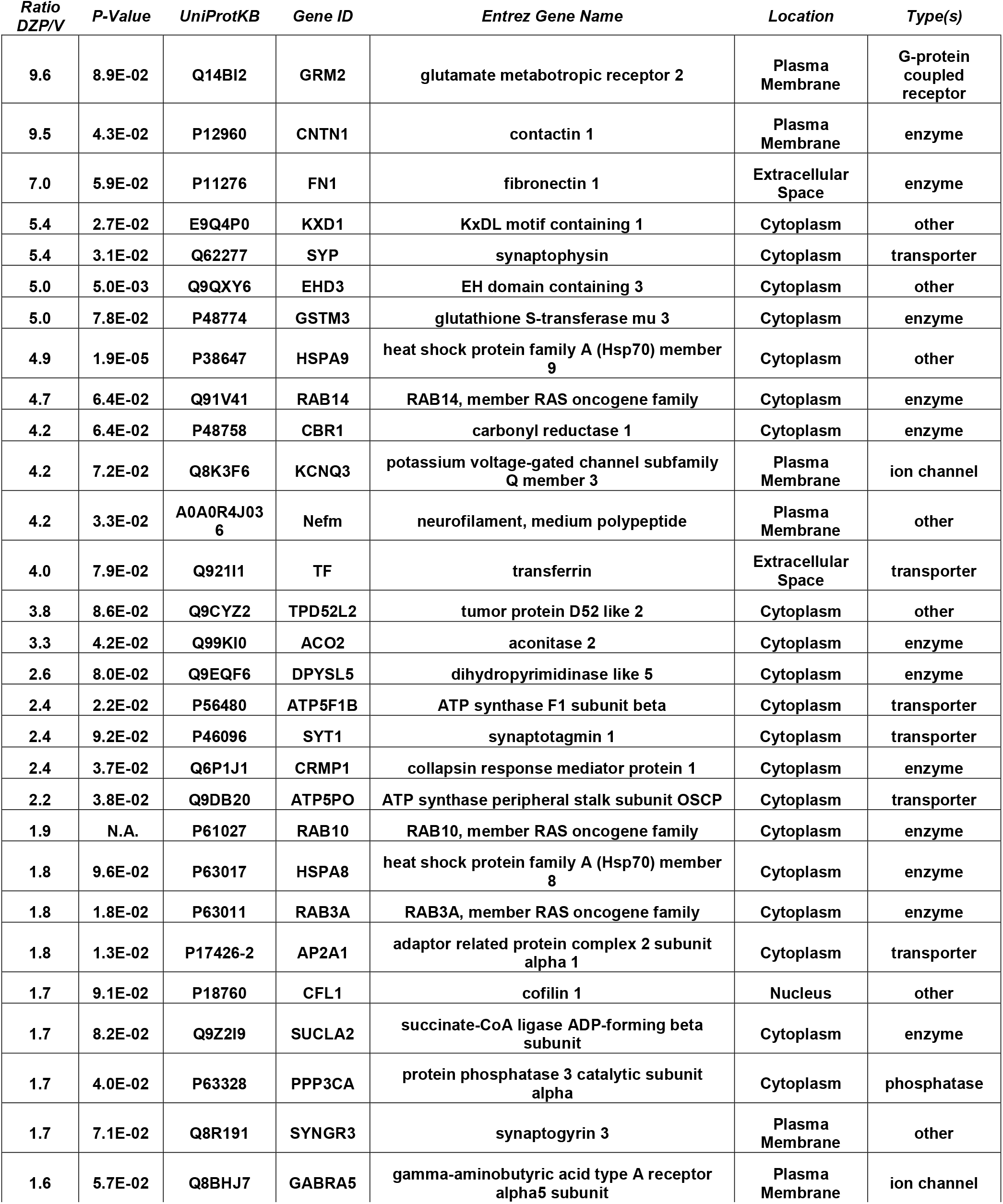
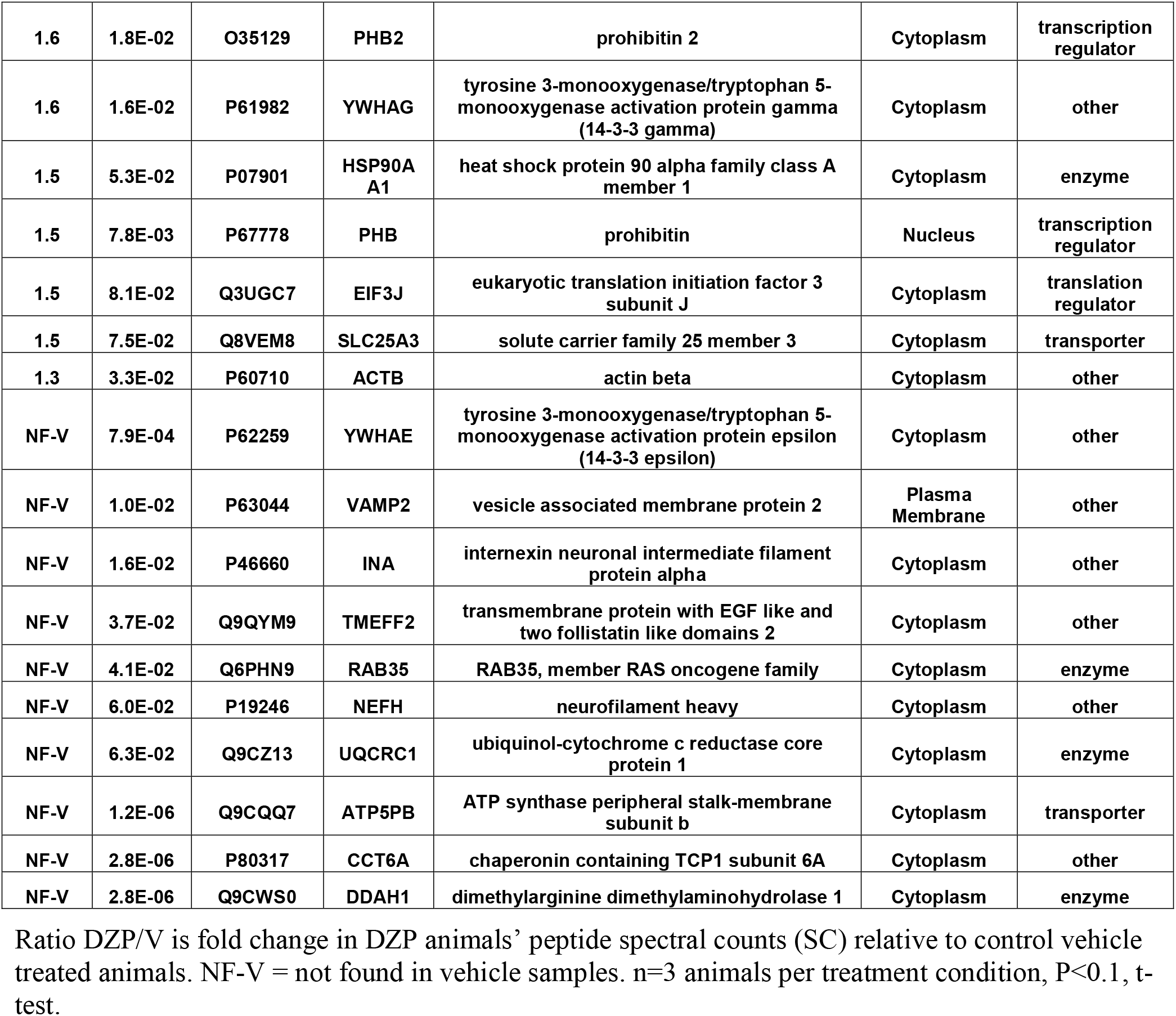
Proteins Demonstrating Increased Association with γ2-GABA_A_Rs after DZP In Vivo by Mass Spectrometry.

**Table 2.**
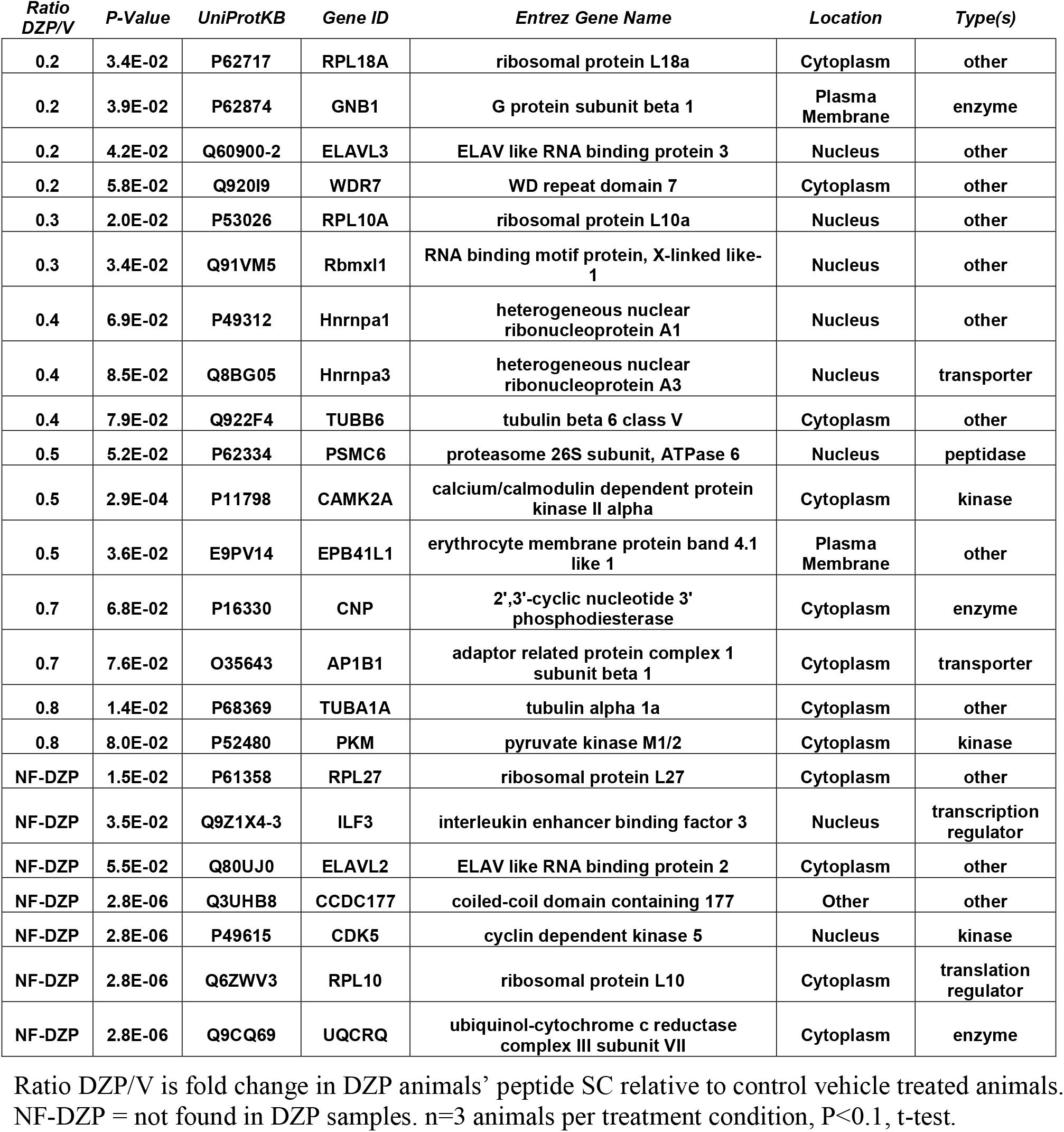
Proteins Demonstrating Decreased Association with γ2-GABA_A_Rs after DZP In Vivo by Mass Spectrometry.

### Bioinformatics Analysis of the γ2 GABA_A_R Interactome

To better understand the consequences of the DZP-induced shift in the γ2-GABA_A_R protein interaction network, protein fold change data was subjected to core Ingenuity Pathway Analysis (IPA). Top enriched canonical pathways with −log(p-value) > 6.2 are shown in Figure 6A. Notably, GABA receptor signaling pathways were highly enriched, as expected, although IPA was unable to determine pathway activation status by z-score analysis. γ2-GABA_A_R association with proteins involved in 14-3-3 mediated signaling and RhoA signaling pathways were greatly increased after DZP (Figure 6A, orange), while interaction with proteins involved in EIF2 signaling and sirtuin signaling pathways were reduced (Figure 6A, blue) relative to vehicle.

**Figure 6.**
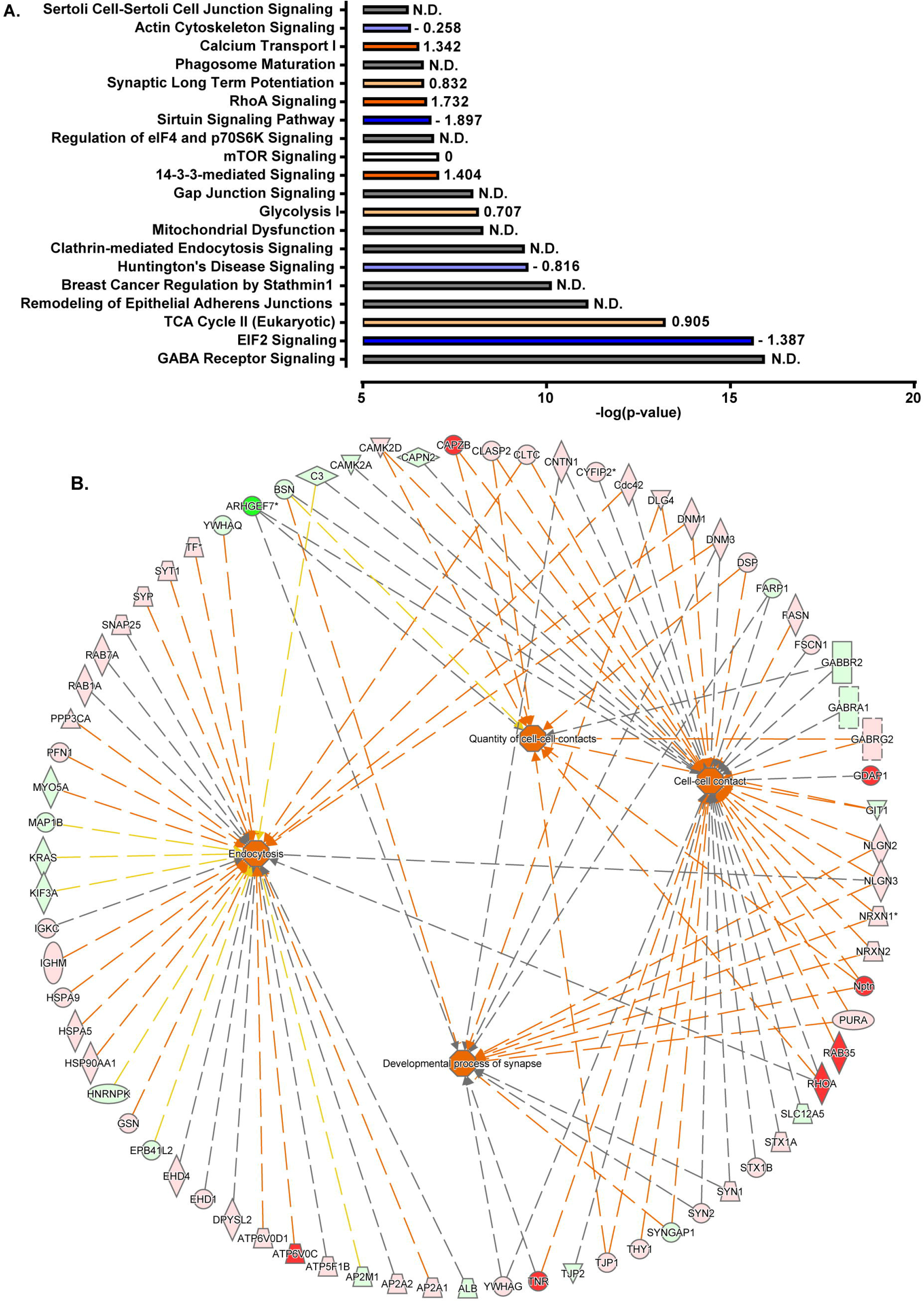
Ingenuity Pathway Analysis Reveals Shifts in Protein Interaction Networks Following DZP Exposure. **(A)** Canonical pathways found to be enriched with γ2-GABA_A_Rs and differentially expressed following DZP administration in vivo. Enriched pathways with −log(p-value) greater than 6.2 were considered as calculated by Fisher’s exact test right-tailed. Values to right of bars represent pathway activation z-score. Positive z-score represents predicted upregulation of a pathway (orange), negative z-score predicts inhibition (blue), z-score = 0 represents no change in pathway (white), while not determined (N.D.) conveys the analysis program was unable to determine a significant change (grey). Intensity of color represents size of z-score value. **(B)** Functional pathway network analysis revealed that DZP treatment resulted in increased γ2-GABA_A_R association with proteins involved in endocytosis, developmental process of the synapse, cell-cell contact, and quantity of cellcell contacts (activation z-score > 2.5). Red = increased measurement, green = decreased measurement, orange = activation of pathway, yellow = findings inconsistent with state of downstream molecule, grey = effect not predicated.

We further examined alterations in functional network association, identifying changes in key trafficking, localization, and cell adhesion pathways in DZP treated mice. Figure 6B lists γ2-GABA_A_R-associated proteins found to be elevated with DZP, contributing to processes such as endocytosis, developmental process of the synapse, cell-cell contact, and quantity of cell-cell contacts (activation z-scores > 2.5). As an additional measurement, we performed gene ontology (GO) database analysis of proteins which were found to be significantly increased (P < 0.1) in DZP treated mice relative to vehicle control (Table 3). GO analysis identified DZP treatment enriched for a number of intracellular trafficking and cellular localization biological pathways that were consistent with the functional network analysis in IPA. Taken together, these results suggest DZP modifies intracellular and surface trafficking of γ2-GABA_A_Rs both in vitro and in vivo.

## Discussion

This work identifies key trafficking pathways involved in GABA_A_R neuroplasticity in response to initial DZP exposure. Using a combination of biochemical and imaging techniques, we identified total γ2 subunit levels are diminished in response to 12-24 h of DZP exposure in vitro and in vivo. Concurrent with the decrease in the overall γ2 pool, we found DZP treatment enhanced ubiquitination of this subunit. Use of an innovative optical sensor for BZD sensitive GABA_A_R (γ2^pH^FAP) in combination with MG dye pulse-labeling approaches revealed DZP exposure moderately enhanced targeting of surface γ2-GABA_A_Rs to lysosomes. Live-imaging experiments with pH 7.4 NH_4_Cl revealed increased intracellular receptor pools, providing further evidence that DZP enhances GABA_A_R lysosomal accumulation, a response reversed by BZD antagonist Ro 15-1788 treatment. We used novel intersubunit FRET based live-imaging to identify that surface synaptic α2/γ2 GABA_A_Rs were specifically decreased after DZP, suggesting these receptor complexes were subjected to ubiquitination, lysosomal targeting, and degradation. In addition to DZP modulation of receptor trafficking, the postsynaptic scaffolding protein gephyrin demonstrated significant plasticity including increased Ser270 phosphorylation and production of gephyrin proteolytic fragments, concurrent with a decrease in total and membrane associated gephyrin levels. Given the fundamental role of gephyrin in scaffolding GABA_A_Rs and regulating synaptic confinement, we used simultaneous FRAP live-imaging of receptors and scaffold in neurons to monitor inhibitory synaptic dynamics. We found ~24 h DZP exposure accelerates both the rate of gephyrin and GABA_A_R exchange at synapses as shown by enhanced fluorescence recovery rates. Control experiments using the BZD antagonist Ro 15-1788 were able to reverse the DZP induced loss of synaptic confinement, reducing gephyrin and GABA_A_R mobility back to vehicle levels. Finally, we used label-free quantitative mass spectrometry and bioinformatics to identify key changes in γ2-GABA_A_R protein association in vivo suggesting enhanced accumulation in cell surface and intracellular trafficking networks. Collectively, this work defines a DZP-induced reduction of gephyrin scaffolding coupled with increased synaptic exchange of gephyrin and GABA_A_Rs. This dynamic flux of GABA_A_Rs between synapses and the extrasynaptic space was associated with enhanced γ2-GABA_A_R accumulation in intracellular vesicles and γ2-GABA_A_R subtype specific lysosomal degradation. We propose DZP treatment alters these key intracellular and surface trafficking pathways ultimately diminishing responsiveness to DZP.

Numerous classical studies have examined gene and protein expression adaptations in GABA_A_R subunits after BZD exposure with minimal agreement that a specific change occurs (Bateson, 2002; Uusi-Oukari and Korpi, 2010; Vinkers and Olivier, 2012). Here molecular mechanistic insight is provided, through direct measurements of enhanced ubiquitination of the γ2 subunit (Figure 2), lysosomal targeting (Figure 3), reduced surface synaptic α2/γ2 GABA_A_R levels (Figure 4), and reduced synaptic confinement (Figure 5) of DZP-sensitive GABA_A_Rs. Together this suggests BZD exposure primarily decreases synaptic retention of γ2 containing GABA_A_R while downregulating surface levels of specific α subunit levels. Ubiquitination of the γ2 subunit by the E3 ligase Ring Finger Protein 34 (RNF 34) (Jin et al., 2014) is the only currently known mechanism targeting internalized synaptic GABA_A_Rs to lysosomes (Arancibia-Carcamo et al., 2009). Due to the requirement of the γ2 subunit in all BZD-sensitive GABA_A_Rs, it is likely that ubiquitination of the γ2 subunit is a contributing factor for increased lysosomal-mediated degradation in response to DZP. Despite a small decrease in the γ2 total protein, changes in surface levels were not significant by biochemical approaches, consistent with evidence that γ2-GABA_A_R surface levels are tightly regulated to maintain baseline inhibition and prevent excitotoxicity. For example, in heterozygous γ2 knockout mice a 50% reduction in γ2 levels appears to be compensated by increased cell surface trafficking, resulting in only approximately a 20% reduction in BZD binding sites in the cortex and a limited reduction in synaptic GABA_A_R clusters. In contrast, homozygous γ2 knockout mice show a complete loss of behavioral drug response to BZD and over 94% of the BZD sites in the brain (GABA binding sites unchanged) and early lethality (Gunther et al., 1995). Similarly, studies have shown that prolonged GABA_A_R agonist or BZD application increases γ2 GABA_A_R internalization in cultured neurons, while surface GABA_A_R levels remain unchanged (Chaumont et al., 2013; Nicholson et al., 2018). Importantly, by using high sensitivity surface GABA_A_R intersubunit FRET measurements we were able to detect a decrease in BZD sensitive α2/γ2 GABA_A_Rs (Figure 4).

The role of inhibitory scaffolding changes in responsiveness to BZD has been largely under investigated. Phosphorylation of gephyrin at Ser270 is mediated by CDK5 and GSK3β, while a partnering and functionally relevant Ser268 site is regulated by ERK1/2 (Battaglia et al., 2018). While the exact signaling mechanism responsible for gephyrin remodeling and phosphorylation in our study is unclear, we have previously shown 30 min treatment with the GABA_A_R agonist muscimol leads to ERK1/2/BDNF signaling, decreased gephyrin synaptic and total levels, and decreased γ2-GABA_A_R at synapses and potentiation by BZDs (Brady et al., 2017). Thus, changes in receptor and scaffold synaptic level and function can occur on the timescale of minutes. Similarly, calpain mediated gephyrin cleavage can occur within 1 minute in hippocampal membranes (Kawasaki et al., 1997), and cleavage products are increased following in vitro ischemia at 30 min and up to 48 hours following ischemic events in vivo (Costa et al., 2015). Additionally, chemically-induced inhibitory long-term potentiation (iLTP) protocols demonstrate gephyrin accumulation occurs concurrent with the synaptic recruitment of GABA_A_Rs within 20 min (Petrini et al., 2014). Collectively, these proteins display a high degree of interdependence across different experimental paradigms of inhibitory synapse plasticity occurring over minutes to days.

A recent work has demonstrated 12 h DZP treatment of organotypic hippocampal slices expressing eGFP-gephyrin caused enhanced gephyrin mobility at synapses and reduced gephyrin cluster size (Vlachos et al., 2013). Here we found the synaptic exchange rate of γ2 GABA_A_Rs and gephyrin to be nearly doubled at synapses in cortical neurons after ~24 h DZP exposure (Figure 4). This effect occurred coincident with the formation of truncated gephyrin cleavage products (Figure 2), which has previously been shown to decrease γ2 synaptic levels (Costa et al., 2015). These findings are also consistent with our previous work showing RNAi gephyrin knockdown doubles the rate of γ2-GABA_A_R turnover at synaptic sites (Jacob et al., 2005). Later quantum dot single particle tracking studies confirmed γ2 synaptic residency time is linked to gephyrin scaffolding levels (Renner et al., 2012). Importantly, GABA_A_R diffusion dynamics also reciprocally regulate gephyrin scaffolding levels (Niwa et al., 2012), suggesting gephyrin and GABA_A_Rs synaptic residency are often functionally coupled. Accordingly, γ2 subunit and gephyrin levels both decrease in responses to other stimuli including status epilepticus (Gonzalez et al., 2013) or prolonged inhibition of IP_3_ receptor-dependent signaling (Bannai et al., 2015).

Increasing receptor synaptic retention enhances synaptic currents, while enhanced receptor diffusion via decreased scaffold interactions reduces synaptic currents. For example, reduction of gephyrin binding by replacement of the α1 GABA_A_R subunit gephyrin binding domain with non gephyrin binding homologous region of the α6 subunit results in faster receptor diffusion rates and a direct reduction in mIPSC amplitude (Mukherjee et al., 2011). Similarly, enhanced diffusion of GABA_A_Rs following estradiol treatment also reduces mIPSCs in cultured neurons and in hippocampal slices (Mukherjee et al., 2017). In contrast, brief DZP exposure (< 1h) reduces GABA_A_R synaptic mobility (Levi et al., 2015) without a change in surface levels (Gouzer et al., 2014), consistent with initial synaptic potentiation of GABA_A_R neurotransmission by DZP. Together with our current findings, this suggests post-translational modifications on GABA_A_R subunits or gephyrin that enhance receptor diffusion are a likely key step leading to functional tolerance to BZD drugs.

It is a significant technical challenge to examine dynamic alterations in receptor trafficking occurring in vivo. To overcome this we examined changes in γ2-GABA_A_R protein association following DZP injection in mice using quantitative proteomics and bioinformatics analysis. This work revealed shifts toward γ2-GABA_A_R association with new protein pathway networks associated with cell surface adhesion, intracellular junctions, synaptic plasticity, endocytosis & recycling and ubiquitination (Figure 6, Table 3), confirming similar fluctuations in membrane and intracellular trafficking occur in vivo and in vitro after DZP. Recent inhibitory synapse proteomics studies have identified a number of new protein synaptic constituents or modulators of GABA_A_R function (Butko et al., 2013; Kang et al., 2014; Nakamura et al., 2016b; Uezu et al., 2016; Ge et al., 2018). We show here that proteins known to have roles in synaptic function and trafficking of membrane receptors show changes in their association with γ2-receptors. For example, the calcium-sensitive kinase CaMKIIα was found to be significantly decreased in interaction with γ2-GABA_A_R following DZP, which can regulate GABA_A_R membrane insertion, synaptic retention and drug binding properties (Churn et al., 2002; Marsden et al., 2010; Saliba et al., 2012; Petrini et al., 2014) (Table 2). Here we report heat shock protein family A (Hsp70) member 8 and heat shock protein family A (Hsp70) member 9 have significantly increased association with γ2 after DZP (Table 1). Overexpression of the heat shock protein family A (Hsp70) member 8 has been shown to reduce gephyrin clustering at inhibitory synapses (Machado et al., 2011), providing a possible mechanistic link between our findings in vitro that DZP diminishes gephyrin scaffolding of γ2-GABA_A_Rs and our in vivo proteomic results. Furthermore, DZP was found to enhance γ2 association with 14-3-3 protein family members (Table 1), which are known mediators of GABA_A_R surface and intracellular trafficking (Qian et al., 2012; Nakamura et al., 2016a). γ2 coassembly with the GABA_A_R α5 subunit was also elevated post DZP exposure (Table 1). Interestingly, the α5 subunit is required for the development of BZD sedative tolerance in mice (van Rijnsoever et al., 2004). Future follow up studies are needed to dissect the individual roles of proteins found to be significantly altered in their association with GABA_A_R, and their physiological and pharmacological importance to BZD tolerance and inhibitory neurotransmission.

Through application of novel and highly sensitive fluorescence imaging approaches combined with in vivo proteomics, we provide unprecedented resolution at both the level of the single neuron and cortex of GABA_A_R synapse plasticity induced by BZDs. Our study reveals that sustained initial DZP treatment diminishes synaptic BZD sensitive GABA_A_R availability through multiple fundamental cellular mechanisms: through reduction of the post-synaptic scaffolding protein gephyrin; shifts toward intracellular trafficking pathways and targeting of receptors for lysosomal degradation; and enhanced synaptic exchange of both gephyrin and GABA_A_Rs. Proteomic and bioinformatics studies using DZP-treated mouse brain tissue provide further evidence that altered γ2-GABA_A_R surface and intracellular trafficking mechanisms play a critical role to the response to DZP in vivo. These results define key events leading to BZD irresponsiveness in initial sustained drug exposure. Future studies utilizing this dual approach will address the neuroadaptations produced by long term BZD use to systematically identify the effects of a critical drug class that has seen a tripling in prescription numbers over the last two decades (Bachhuber et al., 2016).

## Author Contributions

JMLG and TCJ designed the research. JMLG, TCJ, and STW wrote and revised the manuscript. Biochemistry, immunoprecipitation, bioinformatics analysis and fixed and live imaging acquisition and analysis in Figure 1, 3, and 5 was performed by JMLG. FRET imaging and analysis was performed by MJB. Tissue collection was performed by JMLG and SD. Mass spectrometry experiments were designed by TCJ, JMLG and STW. Mass spectrometry was performed by STW and the Weintraub lab and analyzed by JMLG, TCJ, and STW.

## Acknowledgements

This work was supported by funding from National Institutes of Health Grants R56MH114908-01 (to TCJ), T32GM008424 (to JMLG), F31MH117839 (to JMLG), University of Pittsburgh Pharmacology and Chemical Biology Fellowship (to JMLG, William C. deGroat Neuropharmacology Departmental Fellowship), NARSAD young investigator grant (to TCJ) and Pharmacology and Chemical Biology Startup Funds. We thank Jonathan Beckel for technical advice on qRT-PCR and Katarina Vajn for assistance with neuronal cultures. Mass spectrometry analyses were conducted at the UTHSCSA Institutional Mass Spectrometry Laboratory, supported in part by UTHSCSA and the University of Texas System for purchase of the Orbitrap Fusion Lumos mass spectrometer. The expert technical assistance of Sammy Pardo and Dana Molleur is gratefully acknowledged. This manuscript has been released as a Pre-Print at bioRxiv (Lorenz-Guertin et al., 2019).

## Data Availability Statement

The proteomic dataset generated in this study has been submitted to the PRIDE repository [px-submission #318756].

