## Supplementary Data for "Diazepam Accelerates GABA_A_R Synaptic Exchange and Alters Intracellular Trafficking"

Supplementary Material

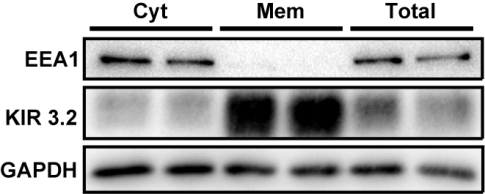

Supplementary Figure 1. Membrane fractionation characterization. DIV 16 neurons were lysed with fractionation buffer and seperated by centrifugation. Western blot analysis shows the intracellular trafficking protein EEA1 is only found in cytosolic and total fractions. Conversely, the plasma membrane protein KIR 3.2 is highly enriched in the membrane fraction.

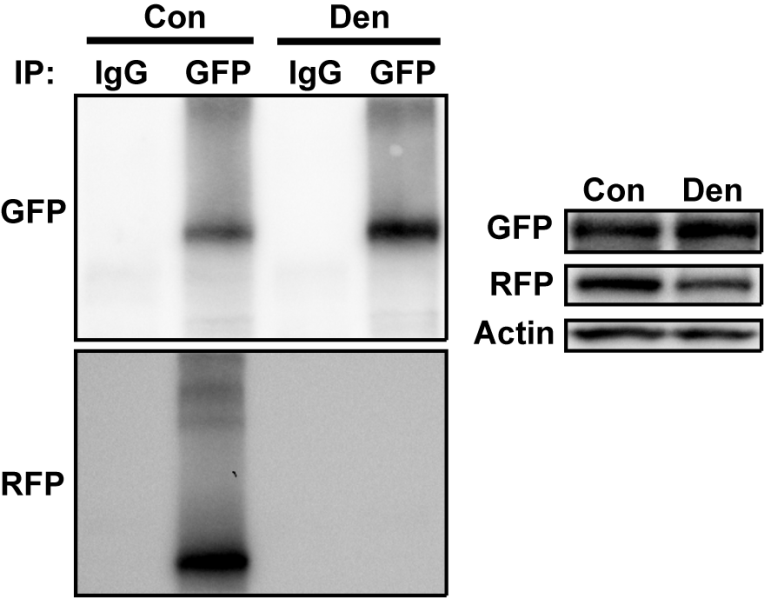

Supplementary Figure 2. Validation of denaturing conditions for γ2 subunit isolation from receptor complex. HEK293 cells were transfected with GABA_A_R subunit DNA γ2^pH^FAP, β3-RFP and α2-HA in a 1:1:1 ratio. Cells were lysed either standard control RIPA conditions (Con) or denaturing conditions (Den) using 1% SDS and 20 min exposure to 50°C heat. Lysates were immunoprecipitated with control IgG or rabbit GFP-antibody. Blots probed with chicken GFP and mouse RFP antibody. Right blots are total lysate expression.

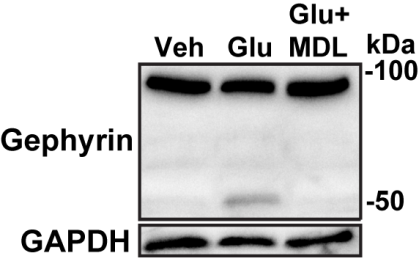

Supplementary Figure 3. Calpain-1 dependent gephyrin cleavage upon glutmate stimulation. DIV 16 neurons were treated with vehicle or 100µM glutamate for 30min in HBS +/- 10µM calpain-1 inhibitor MDL-28170 (MDL). Neurons were then returned to conditioned media for 1.5 h prior to lysis. MDL-28170 treated neurons were in presence of inhibitor throughout experiment. Calpain-1 inhibition mitigates glutamate induced cleavage of gephyrin. Top band indicates full length gephyrin (95 kDa), bottom band indicates cleaved gephyrin (~50 kDA).

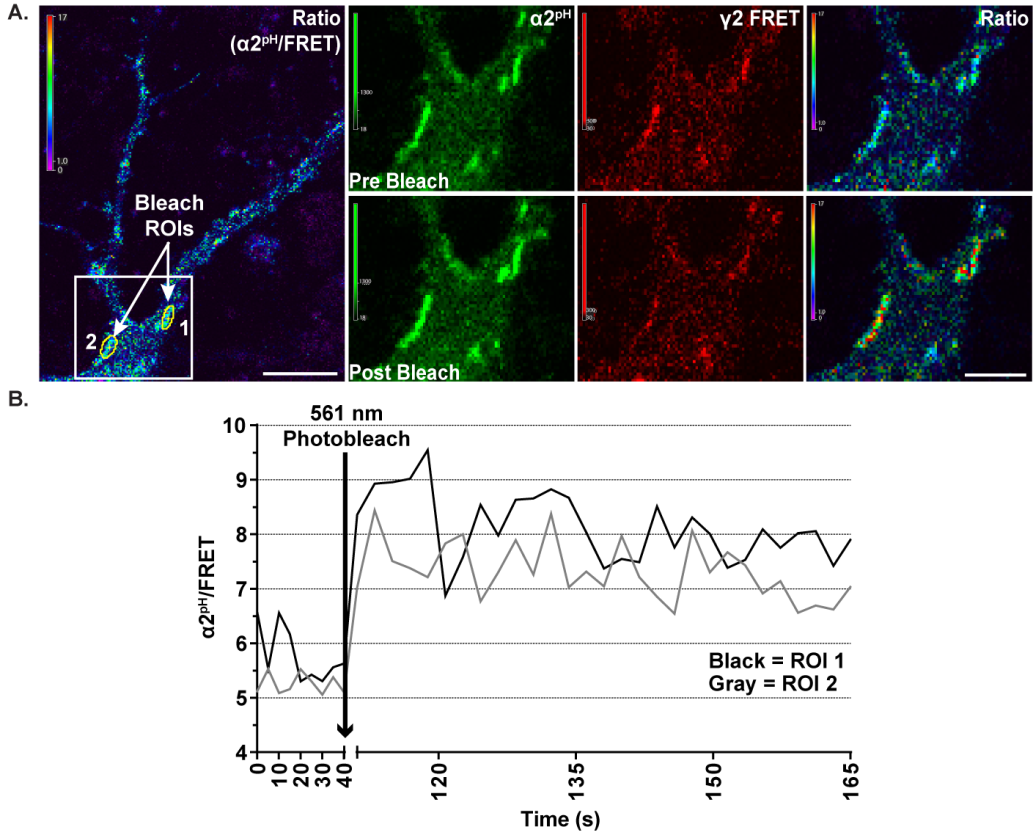

Supplementary Figure 4. α2^pH^ and γ2^RFP^ participate in intermolecular FRET within GABA_A_R complexes. A. FRET occurs between α2^pH^ donor and γ2^RFP^ acceptor in DIV 15-16 cortical neurons. Time-series imaging examined pre bleach and post bleach synaptic sites after photobleaching of γ2^RFP^ (acceptor) with 561 nm laser for 1 minute. B. Acceptor γ2^RFP^ bleaching results in elevated α2^pH^ fluorescence, evidencing previous FRET between fluorophores.

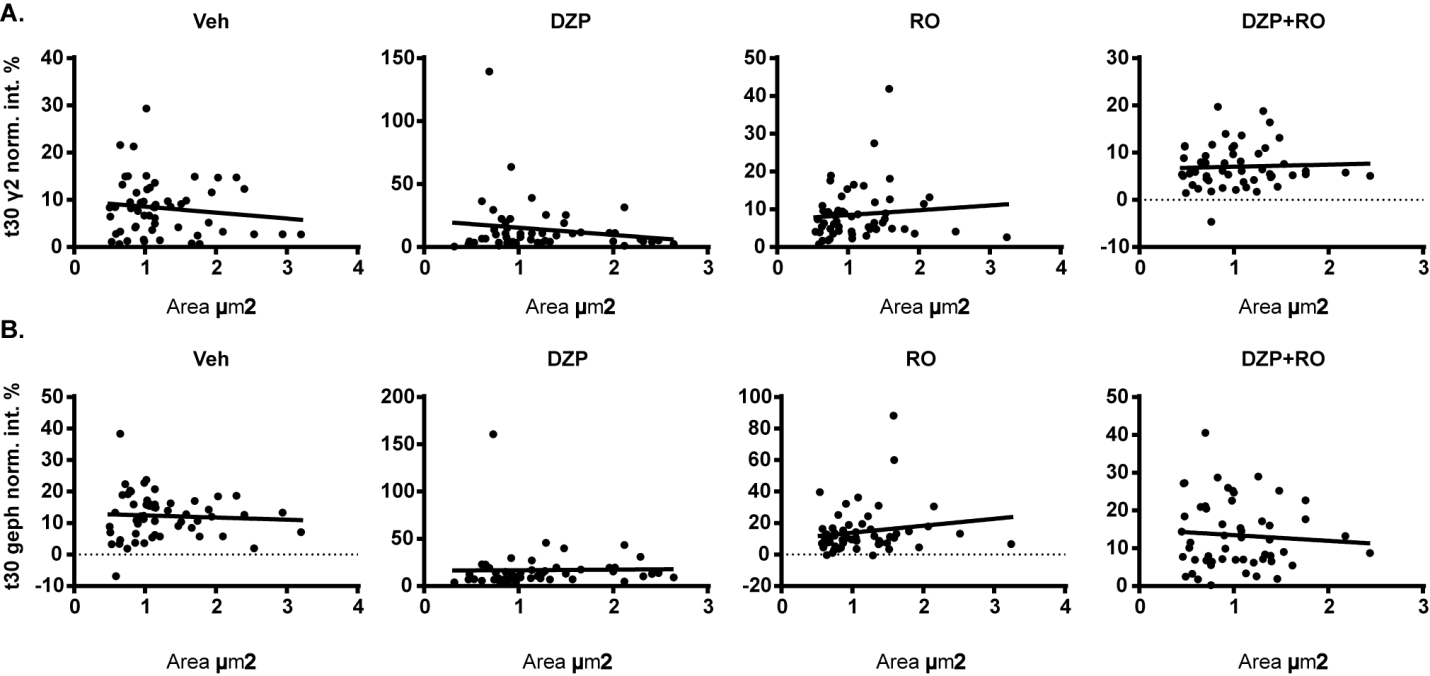

Supplementary Figure 5. Synapse cluster size does not correlate with the rate of fluorescence recovery of gephyrin and γ2. The rate of γ2^pH^FAP **(A)** and RFP-gephyrin **(B)** fluorescence recovery rate at 30 min post-photobleaching was compared to synapse cluster area measured by correlation analysis. There was no significant correlation between fluorescence recovery rate and area across all conditions.

IN EXCEL FILE

Dataset 1. Proteins Identified by Label-Free Mass Spectrometry and Spectral Counts. Identification criteria were: minimum of two peptides; 96% peptide threshold; 1% FDR; 99% protein threshold; identified in at least 3 samples overall or identified 2 of 3 times in a specific treatment group; demonstrated at least 3:1 enrichment over IgG control across at least 3 samples overall.
